# Transient septin sumoylation steers a Fir1-Skt5 protein complex between the split septin ring

**DOI:** 10.1101/2023.01.08.523158

**Authors:** Judith Müller, Monique Furlan, David Settele, Benjamin Grupp, Nils Johnsson

**Affiliations:** Institute of Molecular Genetics and Cell Biology, Department of Biology, Ulm University, James-Franck-Ring N27, D-89081 Ulm, Germany

**Keywords:** diffusion barrier, cytokinesis, chitin synthase, Spa2, polarisome, split-ubiquitin

## Abstract

Ubiquitylation and phosphorylation control composition and architecture of the cell separation machinery in yeast and other eukaryotes. The significance of septin sumoylation on cell separation remained an enigma. Septins form an hourglass structure at the bud neck of yeast cells that transforms into a split septin double ring during mitosis. We discovered that sumoylated septins recruit the cytokinesis checkpoint protein Fir1 to the peripheral side of the septin hourglass. Subsequent de-sumoylation and synchronized binding to the scaffold Spa2 relocate Fir1 in a seamless transition between the split septin rings. Fir1 binds and carries Skt5 on its route to the division plane where the Fir1-Skt5 complex serves as receptor for chitin synthase III. We propose that the opposite positioning of the sumoylated septins and Spa2 creates a tension across the ring that upon de-sumoylation tunnels the membrane-bound Fir1-Skt5 complex through a transiently permeable septin diffusion barrier.

## Introduction

The five mitotic septins of the yeast *Saccharomyces cerevisiae* Cdc11, Cdc12, Cdc10, Cdc3 and Shs1 assemble into hetero-octameric rods that polymerize end to end into apolar filaments (Woods and Gladfelter, 2021; Marquardt et al., 2021). At the beginning of each cell cycle septin filaments form a ring-like structure next to the previous division site through which the directed transport of secretory vesicles creates a new bud that grows and matures into the next daughter cell (Chiou et al., 2017; Pruyne and Bretscher, 2000; Chollet et al., 2020). The septin ring expands later into an hourglass structure in which paired septin filaments are closely aligned at the bud neck along the mother-bud axis (DeMay et al., 2011; Ong et al., 2014). The splitting of the hourglass into two rings prominently marks the beginning of cytokinesis. This transformation involves the loss of the paired filaments with a concomitant formation of new filaments perpendicularly to the mother-bud axis (DeMay et al., 2011; Chen et al., 2020; Vrabioiu and Mitchison, 2006). The contracting actin myosin ring (AMR) subsequently pulls the newly exposed membrane between the two rings towards the center of the bud neck. Chitin synthase II (Chs2) simultaneously synthesizes a primary septum between the two sheets of the constricted plasma membrane. The concerted actions of chitin synthase III (Chs3) and the beta-1,6 Glucan synthase Fks1 layer a secondary septum upon the primary septum. Finally, the daughter cell releases hydrolases that remove the septum to separate both cells (Meitinger and Palani, 2016; Bhavsar-Jog and Bi, 2017; Onishi et al., 2013).

The split septin double ring (SSDR) performs two important functions during cytokinesis. It seals off the membrane of the cytokinesis compartment to keep the membrane-attached proteins within the SSDR, and it serves as a platform from which proteins are sequentially released into the cleavage furrow to take part in the different steps of cell separation (Dobbelaere and Barral, 2004; Meitinger et al., 2013; Meitinger et al., 2011; Schneider, C. et al., 2013; Fang et al., 2010).

The yeast septins Cdc11, Cdc3, and Shs1 are known to be sumoylated during mitosis and de-sumoylated shortly before cell separation (Johnson and Blobel, 1999; Takahashi et al., 1999). Sumoylation involves only a minority of the septins and is restricted to the peripheral region of the septin hourglass facing the mother cell (Johnson and Blobel, 1999). The precise timing of septin sumoylation and de-sumoylation is achieved by the successive export of the SUMO (Smt3) ligase Siz1 and the Smt3 protease Ulp1 from the nucleus into the cytosol (Makhnevych et al., 2007; Takahashi et al., 2001; Johnson and Gupta, 2001; Takahashi et al., 2000). Both enzymes return to the nucleus before a new cell cycle starts (Elmore et al., 2011; Makhnevych et al., 2007; Lewis et al., 2007; Panse et al., 2003). Timing and location imply that septin-sumoylation plays a role during cytokinesis. Although suggestive, the question whether this elaborate modification cycle influences any of the functions of the septins remained open for more than 20 years.

## Results

### Sumoylation recruits Fir1 to the peripheral side of the septin hourglass

Fir1 is predicted to present multiple linear interaction motifs within a predominately unfolded structure (Fig. 1A) (Brace et al., 2019) (Grinhagens et al., 2020; Jumper et al., 2021). Fir1 binds to Smt3 through its SUMO-interacting motif (SIM) and was recently shown to localize to the bud neck during cell abscission (Hannich et al., 2005; Uzunova et al., 2007; Brace et al., 2019; Grinhagens et al., 2020). To test whether binding to Smt3 is required for Fir1’s localization at the bud neck, we compared the cellular distribution of Fir1-GFP to a mutant of Fir1 that lacked its SIM (Fir1_ΔSIM_) through deletion of residues 759-767 and consequently lost its interaction with Smt3 (Fig. 1B). Fir1-GFP appeared at the bud neck 8 to 10 min before the septin ring splits and the AMR starts to contract (Fig 1C). It remained there for approximately 18 min before it rapidly disappeared. Fir1_ΔSIM_-GFP could be detected at the bud neck 10 min later than the wildtype protein, but then showed a similar signal intensity and kinetic of disappearance (Fig. 1C). Fir1-GFP first appeared as a ring at the mother side of the septin structure (Fig. 1D). This ring then shifted between the SSDR during the hourglass-transition and contracted as cytokinesis continued (Fig. 1D) (Brace et al., 2019). Position and timing of the Fir1-ring within the SSDR coincided with the ring-like structure of Fir1_ΔSIM_-GFP when it first appeared at the bud neck (Figs. 1D, S1A). To find out whether septin sumoylation is required for the early appearance of Fir1 at the bud neck (phase I targeting), we measured the fluorescence intensity profiles of Fir1-GFP in cells expressing septins without sumoylatable lysines (*sepΔsumo*), or in cells lacking Siz1 (*siz1*Δ) (Fig. 1E) (Johnson and Gupta, 2001; Johnson and Blobel, 1999). The Fir1-GFP profiles of both mutants were nearly identical, and in their timing very similar to the profile of Fir1_ΔSIM_-GFP. Both lacked phase I targeting (Fig.1E). We conclude that sumoylated septins recruit Fir1 to the periphery of the septin ring shortly before it splits.

**Figure 1:**
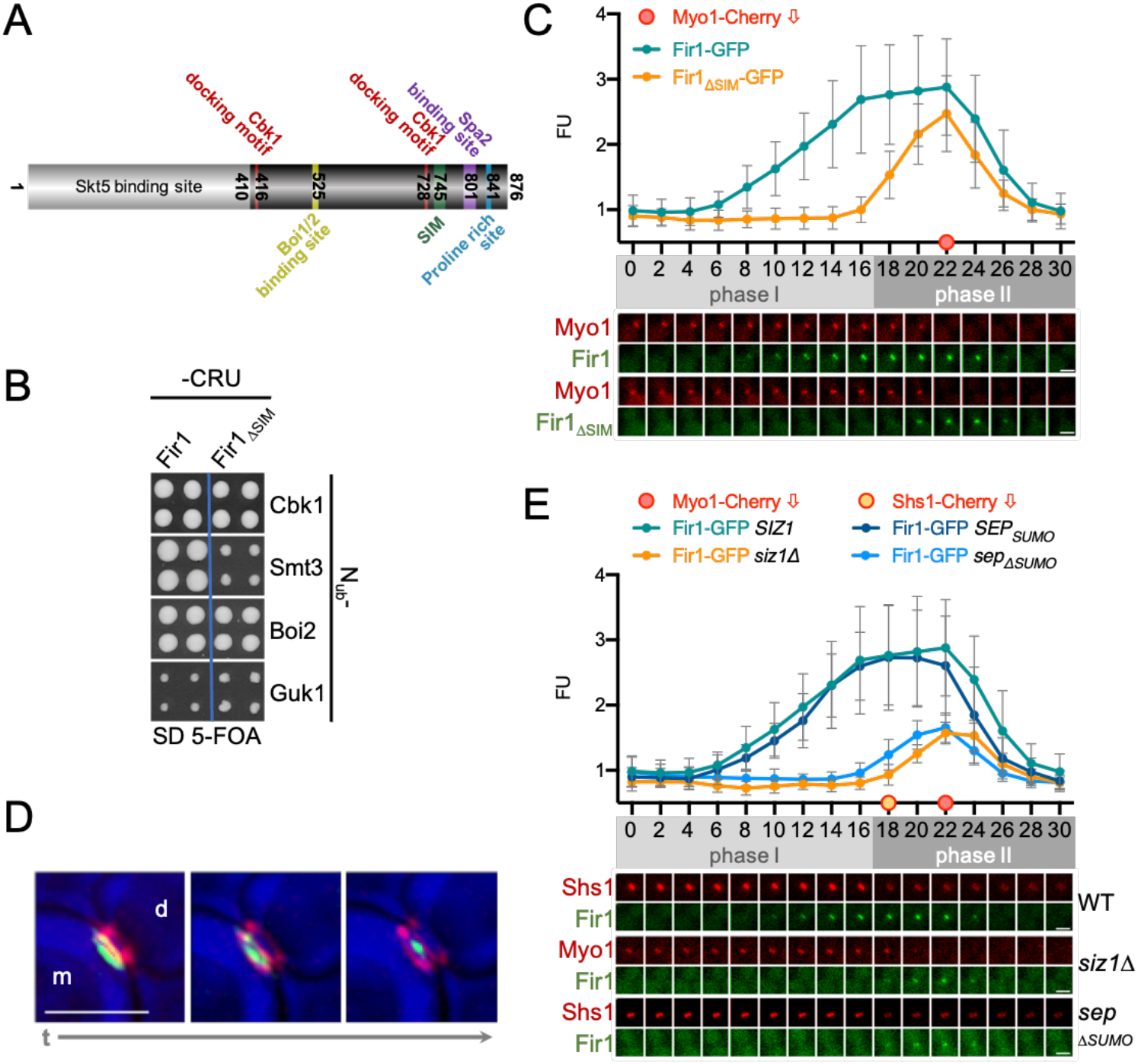
Fir1 is recruited to the bud neck by the sumoylated septins. (A) Cartoon of Fir1 indicating the kno wn and herein discovered binding sites to other proteins. Amino acid positions bordering the motifs are indicated. (B) Cut outs of a Split-Ubiquitin array of diploid yeast cells expressing genomic Fir1CRU (CRU; C-terminal half of Ubiquitin (C_ub_)-R-Ura3) (left panel), Fir1_ΔSIM_CRU (right panel) together with the indicated N_ub_-fusions of known binding partners of Fir1, or the negative control N_ub_-Guk1 (N_ub_, N-terminal half of Ubiquitin). Four independent matings were arrayed as quadruplets on media containing 5-FOA. Colony growth indicates interaction between the fusion proteins. (C) Fluorescence intensities (FU, upper panel) and stills of representative cells (lower panel) coexpressing Fir1-GFP (n=29), or Fir1_ΔSIM_-GFP (n=25) and Myo1-Cherry during cytokinesis. Images were taken every two minutes. Phase I indicates association of Fir1 with the septin ring, phase II localization within the SSDR. The filled red circle indicates the time point of Myo1 disappearance as a result of the final contraction of the AMR and was defined as minute 22 in intensity plots and picture arrays. (D) Deconvoluted images of Fir1-GFP associated with the Cherry-labelled septin ring (Shs1-Cherry) shortly before (left panel), during (middle panel), and shortly after ring splitting (right panel). m and d indicate mother and daughter cell, respectively. (E) As in (C) but with cells coexpressing Myo1-Cherry with Fir1-GFP in wildtype (n=29) or *siz1*Δ-*cells* (n=22), or with cells coexpressing Shs1-Cherry and Fir1-GFP in wildtype (n=19) and *sep*_Δsumo_-cells (n=23). The orange-filled circle at time point 18 indicates the sudden decrease of the Shs1-Cherry signal upon the septin ring split. The significance of the differences between Fir1-GFP signal intensities between wild type and the deletion strains is shown in Fig. S1C. Reintroducing a plasmid-encoded *SIZ1* into the *siz1Δ-cells* restored the targeting of Fir1-GFP (Fig. S1B). All scale bars = 3μm.

During AMR contraction Fir1-GFP moves from its peripheral position between the SSDR (Fig. 1D, S1A; phase II targeting). Neither deleting the SIM in Fir1 nor preventing septin sumoylation delayed phase II targeting of Fir1-GFP (Figs. 1C, E). However, the signal intensities of Fir1-GFP in *siz1*Δ-, or *sep*Δ*sumo-cells* during phase II targeting were substantially reduced when compared to its signal in wildtype cells (Figs. 1E, S1C). These findings argue for a separate receptor that keeps Fir1 at the bud neck after de-sumoylation of the septins. The partial loss of Fir1 from the SSDR in sumoylation-deficient cells might imply that both targeting steps are somehow coupled.

### The polarity scaffold Spa2 attaches Fir1 to the cleavage furrow

We performed a Split-Ub interaction screen to obtain a shortlist of candidate proteins that might anchor Fir1 at the bud neck during phase II targeting (Figs. 2A, S2A, Table S1) (Hruby et al., 2011; Johnsson and Varshavsky, 1994). The polarisome scaffold Spa2 and other subunits of the polarisome were discovered as novel interaction partners of Fir1 (Fig. 2A) (Neller et al., 2015; Tcheperegine et al., 2005; Shih et al., 2005; Dunkler et al., 2021; Valtz and Herskowitz, 1996).

**Figure 2:**
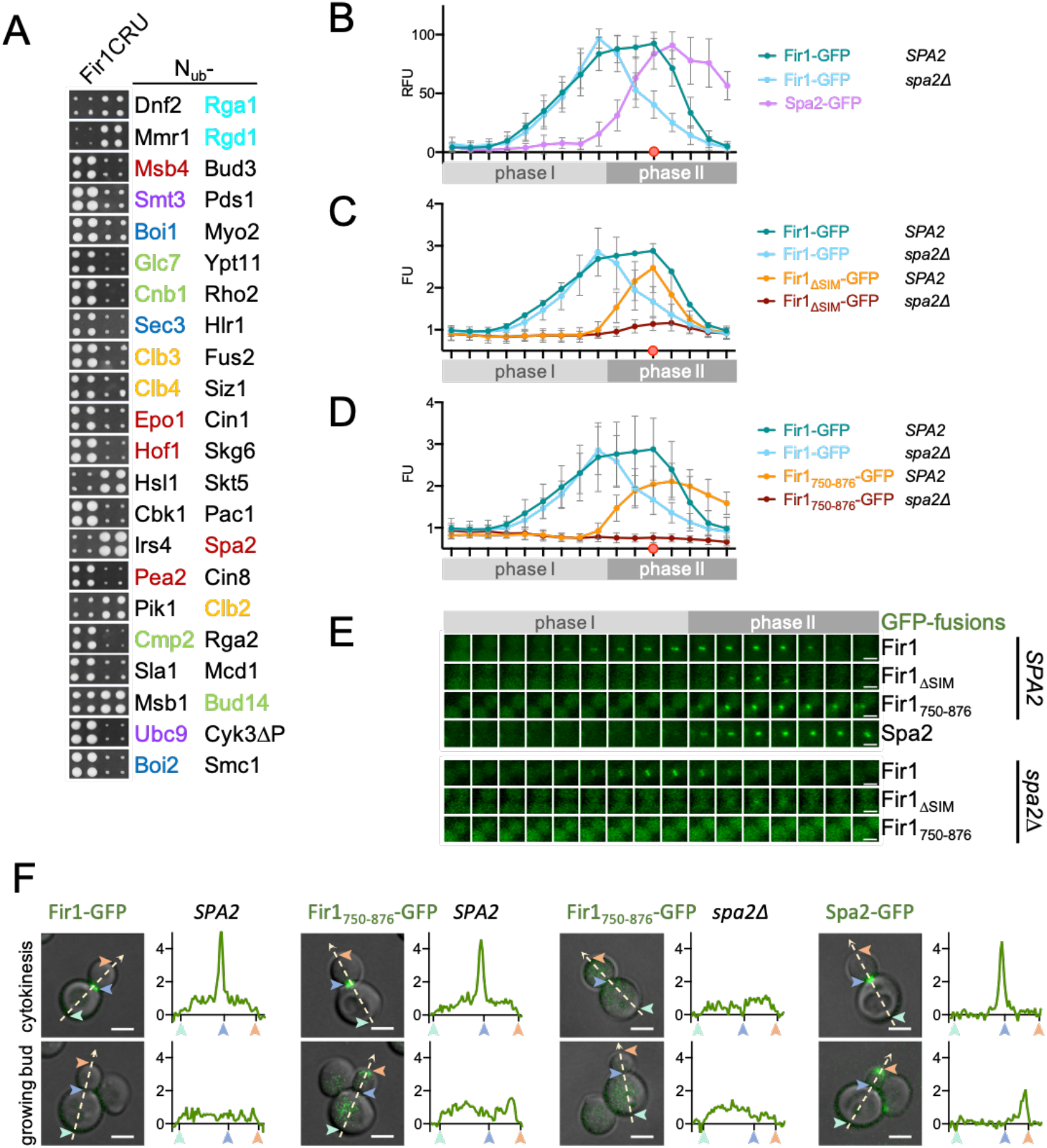
Spa2 recruits Fir1 between the SSDR. (A) Cut outs of a Split-Ub array of 540 diploid yeast cells each expressing genomic Fir1CRU together with a different N_ub_ fusion as in 1A. Identities of the N_ub_-fusions are given next to the respective cut out. Shared colors of the N_ub_ fusions indicate a common process. red: polarisome; purple: SUMO conjugation; dark blue: exocytosis; light blue: Rho GTPase activating protein; yellow: mitotic cyclins; green: phosphatases and their adaptors. The complete analysis is shown in Fig. S2A, and the complete list of binding partners in Table S1. (B) As in (1C) but with cells coexpressing Fir1-GFP with Myo1-Cherry in the presence (n=29) and absence of Spa2 (n=26), or coexpressing Myo1-Cherry with Spa2-GFP (n=18). Intensity values are normalized for each strain by defining the highest intensity value as 100% and the lowest as 0% (RFU). Reintroducing a plasmid-borne *SPA2* into the *spa2*Δ-cells restored the targeting of Fir1-GFP (Fig. S2C). (C) As in (1C) but with cells coexpressing Fir1-GFP or Fir1_ΔSIM_-GFP with Myo1-Cherry in the presence (n=29; n=25) or absence (n=26; n=23) of Spa2. (D) As in (1C) but with cells coexpressing Fir1-GFP or Fir1750-876-GFP with Myo1-Cherry in the presence (n=29; n=22) and absence (n=26; n=20) of Spa2. The significance of the differences between Fir1-GFP and its mutants in wildtype- and *spa2*Δ-strains is shown in Fig. S2E. (E) Stills of representative cells corresponding to the intensity profiles shown in (B) – (D). (F) Fluorescence intensity profiles (shown in arbitrary units) along the polarity axes of cells expressing from left to right Fir1-GFP, Fir1_750-876_-GFP in the presence, or absence of Spa2, or Spa2-GFP. Numbers and arrows correlate positions on the fluorescence intensity profiles with locations in the cells. All scale bars = 3μm.

We considered Spa2 as a candidate receptor as it appeared at the bud neck at about the same time as Fir1_ΔSIM_ during phase II targeting (Figs. 2B, C, E, S2B) (Snyder, 1989). Accordingly, the deletion of Spa2 prevented Fir1-GFP from transferring into the SSDR and nearly abolished the bud neck signal of Fir1_ΔSIM_-GFP during both phases (Figs. 2B, C, E). Reciprocally, the deletion of Fir1 or the absence of septin-sumoylation did not affect the distribution of Spa2 (Fig. S2D). We conclude that the sumoylated septins recruit Fir1 to the bud neck during phase I targeting, whereas Spa2 acts as Fir1 receptor during phase II targeting. A C-terminal fragment of Fir1 (Fir1_750-876_) that starts within the predicted N-terminal beta-strand of the SIM, lacked phase I targeting and recapitulated the dynamic localization of Spa2-GFP at the bud neck (Fig. 2D, E) (Jumper et al., 2021). Similar to Spa2-Cherry, Fir1_750-876_-GFP remained at the site of cell division and later transferred to the tip of the growing bud (Fig. 2F). GFP-fusions of full-length Fir1 and most of its other variants localized to the bud tip at a significantly lower level, almost below the detection limit of our wide-field microscope (Fig. 2F). The deletion of *SPA2* abolished the localization of Fir1_750-876_-GFP equally and completely at bud neck and tip (Figs. 2D, E, F, S2E), thus restricting the potential Spa2-binding site within the C-terminal 126 residues of Fir1.

### Fir1 and Spa2 form a stable protein complex

A stretch within these C-terminal 126 residues of Fir1 is phylogenetically conserved (Fig. 1A). Deleting this region (Fir1_Δ801-820_) specifically abolished the interaction with Spa2, and caused the premature dissociation of Fir1_Δ801-820_-GFP from the bud neck without affecting its initial phase I targeting (Figs. 3A, B, C). The calculated fluorescence intensity profile of Fir1Δ_801-820_-GFP was congruent with the profile of wildtype Fir1-GFP in *spa2*Δ-cells (Figs. 3B, C, E) and complementary to the profile of Fir1_ΔSIM_ (Figs. 3B, C). The simultaneous deletion of the potential Spa2-binding site and the SIM motif completely abolished phase I- and nearly completely phase II targeting of Fir1_ΔSIMΔ801-820_-GFP (Figs. 3B, C).

**Figure 3:**
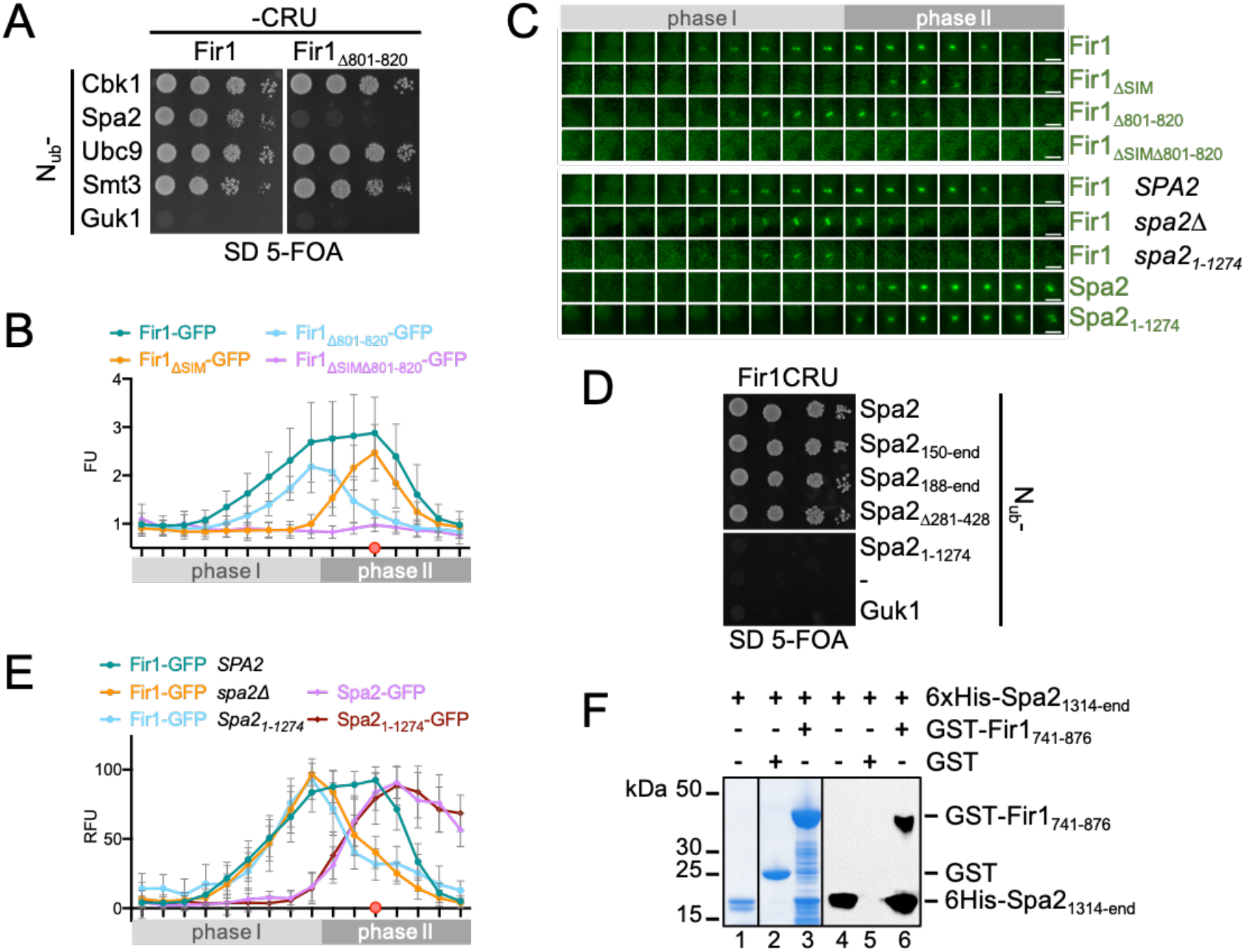
Spa2 is the phase II receptor for Fir1. (A) Split-Ub assay of cells coexpressing Fir1CRU (left panel), or Fir1_Δ801-820_CRU (right panel) with N_ub_-fusions to Spa2 and other binding partners of Fir1, or to Guk1 as negative control. Cells were grown to OD_600_=1 and 4 μl, or 4 μl of a 10-fold serial dilution were spotted on media containing 5-FOA and selecting for the presence of the N_ub_- and Cub fusions. (B) as in (1C) but with cells expressing the GFP fusions to the indicated alleles of *FIR1* (Fir1-GFP n=29; Fir1_ΔSIM_ n=25; Fir1_Δ801-820_ n=23; Fir_ΔSIMΔ801-820_ n=20). (C) Stills of cells corresponding to the intensity profiles shown in (B) and (E). Scale bars = 3μm. (D) Split-Ub assay as in (A) but with cells coexpressing Fir1CRU with the indicated N_ub_ fusion of Spa2, or N_ub_- and N_ub_-Guk1 as non-interacting controls. (E) As in (2B) but with cells coexpressing Myo1-Cherry with Fir1-GFP in wildtype-(n=29), or *spa2*Δ- (n=18), or *spa2_1-1274-_* (n=20) cells, and with cells coexpressing Myo1-Cherry and Spa2-GFP (n=18), or Spa2_1-1274-_ GFP (n=20). (F) Purified 6xHis-Spa_1314-end_ (lanes 1,4) was incubated with GST-Fir1_741-876_ (lane 3, 4) or GST-coupled beads (lanes 2, 5). Glutathione-eluates were separated by SDS-PAGE and stained with Coomassie (lanes 1-3), or transferred onto nitrocellulose, and stained with anti-His antibody (lanes 4-6). Detection of GST-Fir1_741-876_ by the anti-His antibody (lane 6) indicates binding of nitrocellulose-detached 6xHis-Spa_1314-end_ to nitrocellulose fixed GST-Fir1_741-876_ (Fig. S3A).

Testing different fragments of Spa2 in a Split-Ub interaction assay restricted the Fir1 binding site onto Spa2’s C-terminal 192 residues (Fig. 3D). A bacterially expressed GST fusion to Fir1_741-876_ (GST-Fir1_741-876_) precipitated the purified His-tagged C-terminal domain of Spa2 (His-Spa2_1314-end_) (Fig. 3F). We conclude that C-terminal domains of Fir1 and Spa2 form a stable complex that anchors Fir1 between the SSDR of dividing cells. Consequently, cells expressing Spa2 without its C-terminal domain (*spa2_1-1274_*) lacked phase II targeting of Fir1-GFP, while Spa2_1-1274_-GFP still binds indistinguishably to the bud neck (Figs. 3E, C).

### Fir1 is a bud neck receptor for Skt5 during cytokinesis

Fir1 might use its horizontal movement across the septins to carry other proteins piggyback into the space between the SSDR. We measured the fluorescence intensities of GFP-fusions to those Fir1-interaction partners that participate in cytokinesis or abscission, and compared their profiles between wildtype- and *fir1*Δ-*cells* (Figs. 2A, 4). Deleting Fir1 did not affect the distributions of Bud14, Hof1, and Msb1 (Fig. S4A). In contrast, the absence of *FIR1* reduced or abolished the phase I targeting of its binding partners Cbk1, Boi1, and Skt5 (Figs. 4A-C, F, S4B, S4C) (Brace et al., 2019; Grinhagens et al., 2020). While the amounts of Cbk1 and Boi1 recovered to wildtype-like levels during phase II targeting, the concentration of Skt5 in the SSDR was significantly reduced (Figs. 4A-C, S4C). Contrary to the other interaction partners, Skt5 seems to require the continuous binding to Fir1 for its two-step translocation into the SSDR.

**Figure 4:**
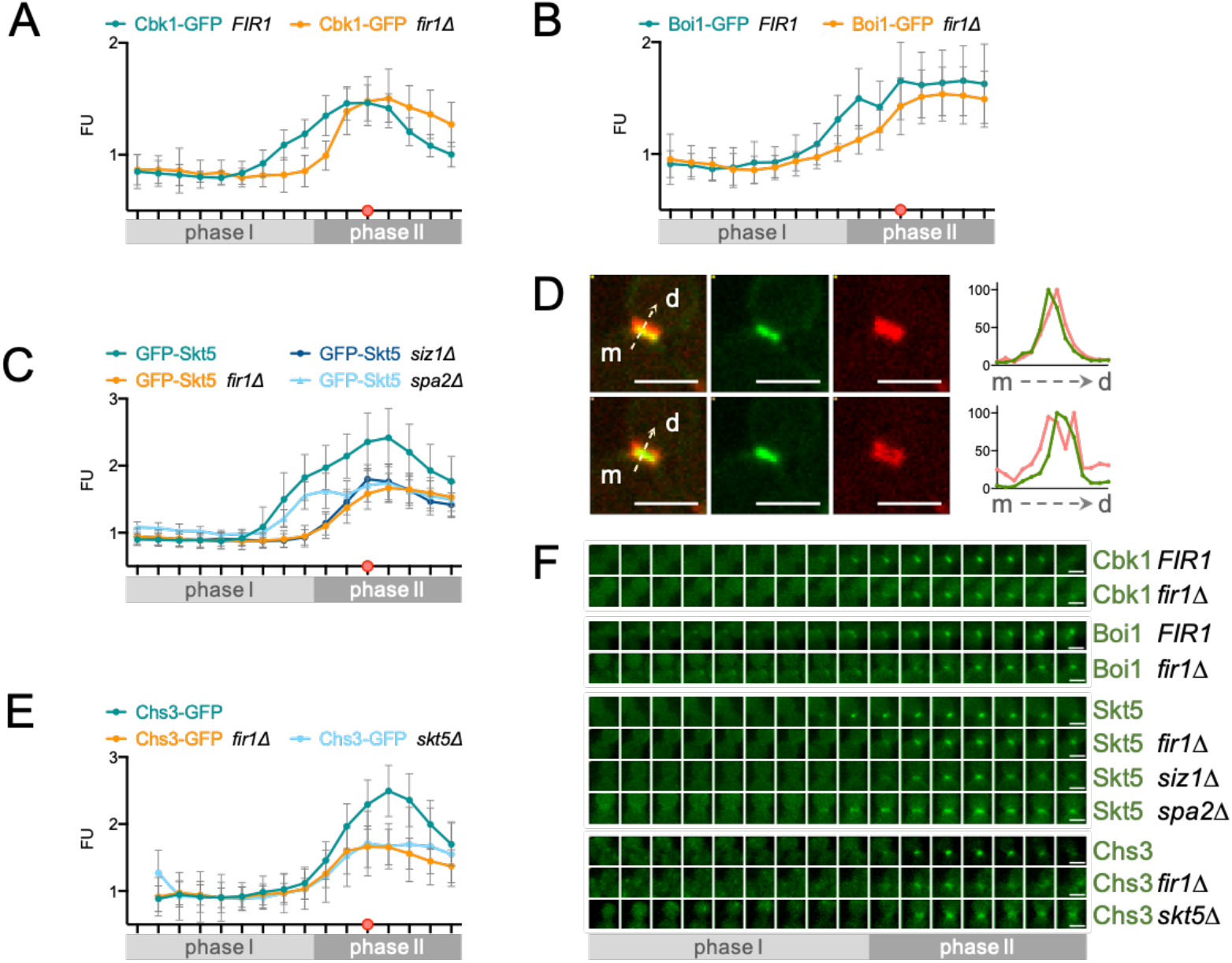
Fir1 tunnels Skt5 between the SSDR. (A) As in (1C) but with cells coexpressing Cbk1-GFP together with Myo1-Cherry in the presence (n=20) or absence (n=20) of *FIR1*. (B) As in (1C) but with cells coexpressing Boi1-GFP with Myo1-Cherry in the presence (n=20) or absence (n=25) of *FIR1*. (C) As in (1C) but with cells coexpressing Skt5-GFP with Myo1-Cherry in the presence (n=22) or absence of *FIR1* (n=27), of *SIZ1* (n=23), or of *SPA2* (n=21). (D) Images of GFP-Skt5 associated with the Cherry-labelled septin ring shortly before ring splitting (upper panels from left to right: overlay, GFP-, Cherry-channel), and shortly after ring splitting (lower panels from left to right: overlay, GFP-, Cherry-channel). m and d indicate mother and daughter cell respectively. Intensity plots of the GFP (green)- and Cherry (red)-channels along the arrows orthogonal to the septin rings are shown in the upper and lower right panels as relative fluorescent units. (E) As in (1C) but with cells coexpressing Chs3-GFP with Myo1-Cherry in wildtype (n=20), and in the absence of *FIR1* (n=20), or *SKT5* (n=20). The significance of the differences between the profiles in Fig. 4B, C, E are shown in Fig. S4B-D. (F) Stills of cells and corresponding to the intensity profiles shown in (A) – (C). Scale bar = 3μm.

Skt5 activates Chs3 and thus determines where and when chitin synthesis occurs (Ono et al., 2000; DeMarini et al., 1997). Recruited by the Bni4-Glc7 complex, Skt5 forms a ring around the incipient bud site and the neck of small budded cells (Larson et al., 2008; DeMarini et al., 1997; Kozubowski et al., 2003). It subsequently spreads out on the plasma membranes of mother- and daughter cell before being concentrated independently of Bni1-Glc7 at the bud neck during mitosis (DeMarini et al., 1997; Gohlke et al., 2018; Kozubowski et al., 2003). Microscopy of GFP-Skt5 expressing yeast cells revealed that Skt5, similar to Fir1, contacted the septin ring first at the mother side before entering the space between the SSDR (Fig. 4D). Inhibiting Smt3 modification of the septins in a *siz1Δ*-strain abolished phase I targeting of Skt5 and decreased the amount of GFP-Skt5 that transfers between the SSDR during phase II targeting (Figs. 4C, F, S4C). The deletion of *SPA2* did not influence the targeting of Skt5 to the septin hourglass but substantially reduced its amount between the SSDR (Figs. 4C, F, S4C). These experiments confirm Fir1 as the mitosis-specific bud neck receptor for Skt5.

### Fir1-Skt5 supports the targeting of Chs3 to the SSDR

Skt5 activates Chs3 at the bud neck of dividing cells (DeMarini et al., 1997; Kozubowski et al., 2003). The bud neck appearance of Chs3 coincided with the phase II targeting of the Fir1-Skt5 complex (Fig. 4E) (Okada et al., 2020). By comparing the fluorescence intensity profiles between wildtype- and *fir1Δ-cells* we could show that *fir1*Δ-*cells* enriched significantly less Chs3-GFP at the bud neck than wildtype cells (Figs. 4E, F, S4D). Chs3-GFP also stayed less focused within the space between the SSDR of *fir1*Δ-*cells* (Fig. 4F). The deletion of Skt5 reduced the Chs3-GFP signal to a similar extent (Figs. 4E, F, S4D). We conclude that the Fir1-Skt5 complex supports the targeting and/or anchorage of Chs3 during cellular abscission (Reyes et al., 2007).

### Fir1 contains a separate binding site for Skt5

Skt5 is attached to the plasma membrane by a C-terminally coupled farnesyl moiety (Grabinska et al., 2007). Our findings suggest that Fir1 first links Skt5 to the septin hourglass before the complex glides on the membrane across the septin diffusion barrier between the SSDR. This model implies that the interface of Fir1 for Skt5 should not overlap with its interfaces for Smt3 and Spa2. Indeed, Split-Ub interaction analysis of fragments of Fir1 localized the binding site for Skt5 to the supposedly disordered N-terminal 410 residues of the protein (Figs. 5A, 1A) (Jumper et al., 2021). A GST-fusion to the N-terminal region of Fir1 (GST-Fir_11-410_) precipitated the bacterial expressed His-Skt5 and a fragment of Skt5 that harbored its central SEL domain (His-Skt5_146-550_) (Fig. 5B). The Fir1-Skt5 complex could also be reconstituted by incubating nitrocellulose membrane-tethered GST-Fir1_1-410_ with *E.coli-extracts* containing His-Skt5_146-550_ (Fig. 5C). We conclude that the interfaces of the Fir1-Skt5 complex are provided by the central SEL-domain of Skt5 and the N-terminal half of Fir1.

**Figure 5:**
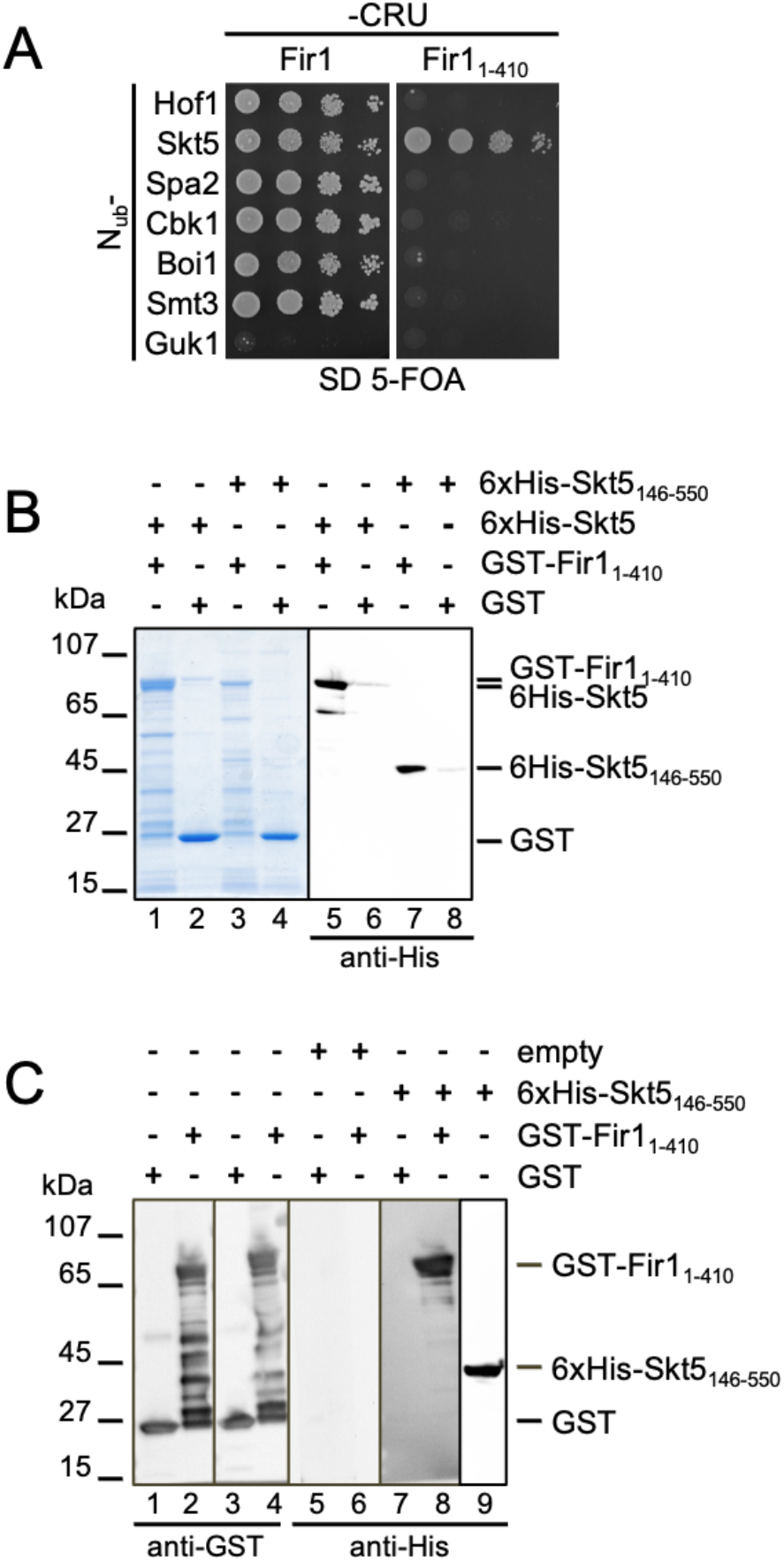
Skt5 and Fir1 form a protein complex. (A) Split-Ub assay as in Fig. 3A but with cells coexpressing Fir1CRU (left panel) or Fir1_1-410_CRU (right panel) with the indicated N_ub_ fusions of binding partners of Fir1, or N_ub_-Guk1 as a negative control. (B) Extracts of *E.coli* expressing 6xHis-Skt5 (lanes 1, 2, 5, 6), or 6xHis-Skt5_146-550_ (lanes 3, 4, 7, 8) were incubated with GST-Fir1_1-410-_ (lane 1, 3, 5, 7), or GST-coupled beads (lanes 2, 4, 6, 8). Glutathione-eluates were separated by SDS-PAGE and stained with Coomassie (lanes 1-4), or transferred onto nitrocellulose, and stained with anti-His antibody (lanes 5-8). Input fractions are shown in Fig. S5A. (C) Extracts of *E.coli* cells expressing GST-Fir11-410 (lanes 2, 4, 6, 8), GST (lanes1, 3, 5, 7), or 6xHis-Skt5_146-550_ (lane 9) were separated by SDS PAGE, transferred on nitrocellulose and either probed directly with anti-GST antibody (lanes 1-4), or anti-His antibody (lane 9), or incubated first with diluted extracts of *E.coli* cells expressing no additional protein (lanes 5, 6), or 6xHis-Skt5146-550 (lanes 7, 8), before being incubated with anti-His antibody (lanes 5-8).

### Degradation of Fir1 resets the targeting of Skt5

Fir1 is a substrate of the Cdh1-activated ubiquitin ligase APC and becomes degraded at the end of cytokinesis (Ostapenko et al., 2012). Accordingly, deletion of Cdh1 raised cytosolic levels of Fir1 and lead to prominent tip-staining of Fir1-GFP during bud growth (Figs. 6A, B, C). Beside a strong cytosolic signal, the fluorescence intensity profile of Fir1-GFP in *cdh1*Δ cells showed phase I targeting and a stronger and longer enrichment between the SSDR (Figs. 6A, B, C, S6A). Deviating from Fir1, GFP-Skt5 lacked phase I targeting in *cdh1*Δ-*cells* and showed a reduced enrichment between the SSDR (Figs. 6B, D, S6B). We conclude that the degradation of Fir1 at the end of cytokinesis is required to properly reset the mechanism for Skt5 enrichment in the next cell cycle.

**Figure 6:**
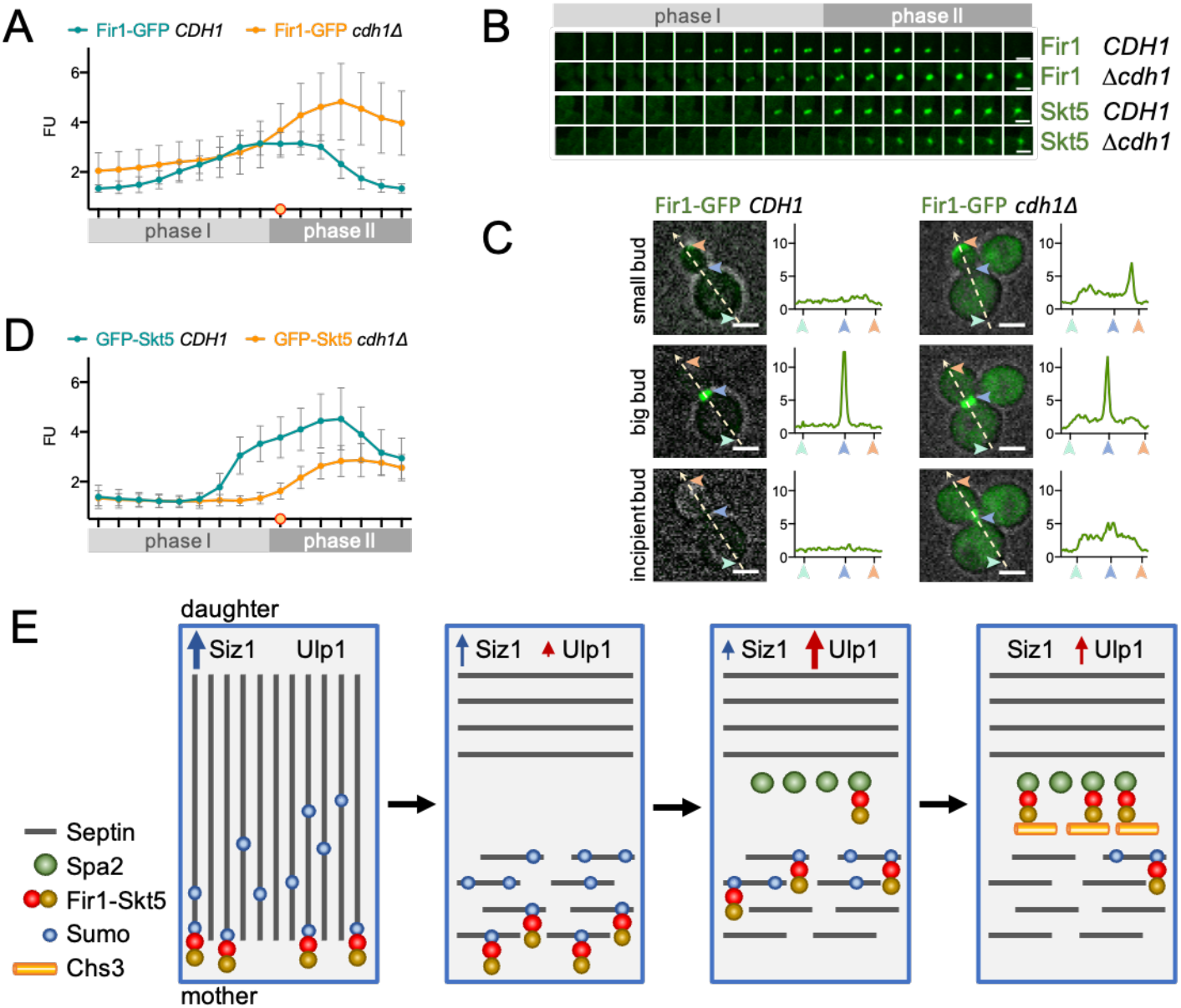
(A) As in (1C) but with cells coexpressing Fir1-GFP together with Shs1-Cherry in the presence (n=24) or absence of *CDH1* (n=25). The fluorescence intensities (FU) are shown as ratio to extracellular background rather than intracellular control due to high cytosolic Fir1-GFP signal in *cdh1Δ-cells* (see Materials and Methods). (B) Stills of cells corresponding to the intensity profiles shown in (A) and (D). (C) Fir1-GFP fluorescence intensity profiles (arbitrary fluorescent units) along the polarity axis in a wildtype cell (left panels), and a *cdh1*Δ-*cell* (right panels). Shown from top to bottom: cells during S-phase, during cytokinesis, and during G1-phase. Numbers and arrows correlate positions on the fluorescence intensity profiles with locations in the cells. (D) As in (1C) but with cells coexpressing GFP-Skt5 and Shs1-Cherry in the presence (n=23) or absence (n=25) of *CDH1*. The orange-filled circle indicates splitting of the septin ring in (A) and (D). Scale bars = 3μm. The significance of the differences between the profiles in Figs.6A, D are shown in Figs.S6A, B. (E) A model how the Fir1-Skt5 complex transfers into the space between the split septin rings. From left to right: Sumoylation at the mother side of the intact septin hourglass concentrates the membrane-bound Fir1-Skt5 complex at the edge of the hourglass. The transformation into the split septin rings reduces the number and length of the septin filaments and switches the direction of the remaining filaments perpendicular to the mother-daughter axis. This process creates temporal holes within the septin grid (only shown for the mother side). Fir1-Skt5 diffuses through these holes along the unoccupied Smt3-sites. The Spa2-trap within the split septin rings and the gradual de-sumoylation from the mother side direct the diffusion of the Fir1-Skt5 complex into the space between the SSDR. The holes are filled by repolymerization, and Chs3 is finally captured and activated by the Skt5-Fir1-Spa2 complex.

## Discussion

Fir1 is the first identified sumoylation-dependent binding partner of the septins. In line with the kinetics of their sumoylation, Fir1 is recruited to the septins shortly before the onset of cytokinesis and is released as soon as the transition of the hourglass to the SSDR occurs. The released Fir1 is retained at the bud neck exclusively by the perfectly scheduled appearance of its newly discovered binding partner Spa2. Sequential interactions move Fir1 from the periphery of the septin hourglass to the center of the SSDR. Intuitively, crossing the septin barrier should proceed in two steps: Release from the de-sumoylated septins into the cytosol, followed by binding to the SSDR-located Spa2. However, Fir1’s binding partner Skt5 is attached to the membrane and might prevent the complex from freely entering the cytosol after septin de-sumoylation (Meissner et al., 2010; Larson et al., 2008; Reyes et al., 2007). As neither GFP-Skt5 nor Fir1-GFP have ever been detected in endosomal compartments during cytokinesis, we suggest that the Fir1-Skt5 complex tunnels straight across the septin barrier without a detour through the secretory system. As Skt5 and Chs3 move into the SSDR *via* different routes this mechanism would avoid that Skt5 prematurely activates Chs3.

In this unusual transfer mechanism septin sumoylation first recruits Skt5 to the edge of the septin hourglass (Fig. 6E). Upon the hourglass to double-ring transition, the septin-gate opens for a brief moment and the Fir1-Skt5 complexes follow their concentration gradient by diffusion along the sumoylated septins toward the center. Gradual de-sumoylation by Ulp1 from the mother side of the ring impedes backward movement, while Spa2 traps the arriving Fir1-Skt5 complexes within the double ring.

We observed that the absence of sumoylation seems to affect the accumulation of Fir1 between the SSDR more severely than the loss of the Sumo binding site in Fir1_ΔSIM_ (Figs. 1C, E). This finding might point to an additional role of septin sumoylation on the state of the septins during the hourglass-ring transitions or its properties as diffusion barrier. These features might correspond with findings in mammalian cells, where sumoylation was shown to alter the bundling properties of the septins during cytokinesis (Ribet et al., 2017).

There is no direct experimental evidence that membrane-attached proteins diffuse across the septin barrier during the hourglass-SSDR transition. However, this transition is accompanied by the disappearance of the majority of the septin filaments. The remaining septin units show an increased mobility, and temporarily lose their uniform orientation (DeMay et al., 2011; Dobbelaere et al., 2003; Ong et al., 2014). One of the proposed models to explain this behavior postulates a depolymerization of the paired filaments followed by repolymerization into circumferential filaments. This is not only compatible with the data but also with a temporal break down of the diffusion barrier (DeMay et al., 2011; Ewers, 2011).

A permeable septin gate alone should already allow a significant fraction of Skt5 to passively diffuse into the SSDR. Incoming interaction partners like Chs3, or Hof1 might then trap the protein between the SSDR (Oh et al., 2017; DeMarini et al., 1997; Reyes et al., 2007). In agreement with this prediction, we observed that the loss of sumoylated septins, or the deletion of either Fir1 or Spa2, did not completely prevent Skt5 from accumulating between the SSDR (Fig. 4C). This Fir1-independent enrichment at the bud neck could also explain why Fir1-, Fir1_ΔSIM_-, or Fir1_Δ801-820_-GFP, all retaining their Skt5 binding site, still show some residual accumulation between the SSDR of *spa2*Δ cells, whereas Fir1750-876-GFP does not (Figs. 2, 3).

Like other cytokinesis-relevant proteins, Fir1 is degraded by Cdh1-activated APC at the end of the cell cycle (Ostapenko et al., 2012; Tully et al., 2009). Impairment of this degradation causes Fir1, that is partly bound to Spa2, to accumulate in the cytosol and to compete with the Fir1-Skt5 complex for the limited number of sumoylated septins. As a consequence, less Skt5 is recruited to the septins and enriched between the SSDR. Our experiments show that the cell tightly controls Fir1 levels to efficiently concentrate the Fir1-Skt5 complex between the SSDR which in turn enriches and focuses Chs3 at the plane of cell separation (Fig. 4E). The resolution of our experiments cannot distinguish whether Fir1-Skt5 promotes exocytosis or prevents endocytosis of Chs3. The binding to Spa2 however brings Fir1-Skt5 in close proximity to the polarisome member Msb4 (Fig 2A). As Rab-GAP, Msb4 stimulates the hydrolysis of Sec4_GTP_ and drives the subsequent attachment of vesicles to the plasma membrane (Donovan and Bretscher, 2015). By simultaneously binding to Chs3, Spa2, Sec3, and the plasma membrane, the Skt5-Fir1 complex could position Chs3-containing vesicles close to the polarisome and the constricting membrane and might thus accelerate their fusion (Fig. 2A, Table S1).

Fir1 was recently discovered as a central component of a new cytokinesis checkpoint that ensures that cell separation is only initiated once the preceding steps have been successfully performed (Brace et al., 2019). The execution of this checkpoint can be provoked by deleting Chs2 or proteins like Cyk3 or Inn1, that stimulate the chitin synthesis of the primary septum (Brace et al., 2019; Foltman et al., 2016; Devrekanli et al., 2012). The additional lack of Fir1 impairs the growth of these cells and leads to a much thinner secondary septum with a reduced chitin content (Brace et al., 2019). Our study identifies Fir1 as the mitosis-specific carrier of the activator of Chs3 and thus offers a molecular explanation for these observations: The loss of Fir1 reduces the amount of Skt5 at the bud neck, possibly below the level that can still compensate for impaired primary septum formation in *chs2*Δ-, *inn1*Δ-, or *cyk3*Δ-cells. Fir1 might thus respond to defects in cytokinesis by two mechanisms: Attenuating the progress into cell separation while simultaneously promoting a rescue pathway.

## Material and Methods

### Growth conditions, cultivation of yeast strains and genetic methods

All yeast strains were derivatives of JD47, a segregant from a cross of the strains YPH500 and BBY45, and are listed in Table S2 (Dohmen et al., 1995). Yeast strains were cultivated in SD or YPD media at the indicated temperatures. Media preparation followed standard protocols (Glomb et al., 2020). SD medium for Split-Ub assays contained in addition 1 mg/ml 5-fluoro-orotic acid (5-FOA, Formedium, Hunstanton, Norfolk, UK). Gene deletions and promoter replacements by *P_MET17_* were performed by homologous integration of the cassettes derived by PCR from the plasmids pFA6a-hphNT1, pFA6a-natNT2, pFA6a-kanMX6, pFA6a-CmLEU2 or pYM-N35 (Bähler et al., 1998; Janke et al., 2004). *E.coli* XL1 blue cells were used for plasmid amplification and grown at 37°C in LB medium containing antibiotics. *E.coli* BL21 cells were used for protein production and were grown in LB or SB medium at 37°C.

### Generation of plasmids and yeast strains

Detailed lists of all plasmids used in this study are provided in Table S3. Genomic gene fusions were obtained as described (Wittke et al., 1999; Neller et al., 2015; Dunkler et al., 2012). In brief, fusions of *GFP, mCHERRY*, or *CRU* to *SPA2, MSB1, CBK1, HOF1, BUD14, CHS3, BOI1, SHS1, MYO1 and in some cases to FIR1*, were constructed by PCR amplification of the respective C-terminal ORFs without stop codon from genomic DNA. The obtained DNA fragments were cloned via *EagI* and *Sal*I restriction sites in front of the *CRU-, GFP-, mCherry* -module on a pRS303, pRS304 or pRS306 vector (Wittke et al., 1999). The plasmids were linearized using a single restriction site within the C-terminal genomic DNA sequence and transformed into yeast. Successful integration was verified by PCR of single yeast colonies with diagnostic primer combinations using a forward primer annealing in the target ORF but upstream of the linearization site, and a reverse primer annealing in the C-terminal module. Most *Fir1-* fusions were created by introducing a PCR fragment carrying GFP and a *hphNT1* selection cassette flanked by 45bp homologous to the 3’-end of the *FIR1* open reading frame and the 5’-end of its 3’ non-translated region, respectively. *GFP-SKT5* was generated *via* CRISPR/Cas9-mediated insertion of a PCR-fragment encoding GFP flanked by 45bp homologous to the region upstream of the start codon of *SKT5* and the 5’-end of its open reading frame. Deletions and base exchanges in the genomic copies of *FIR1*, and *SPA2* were achieved by CRISPR/Cas9 manipulation using plasmid pML104 or pML107 containing specific 20mer guide-RNA sequences, and template oligonucleotides for exchanging the information on the genomic DNA (Table S4) (Laughery et al., 2015). The integrity of all integrated PCR-fragments and introduced mutations was verified by PCR-amplification and sequencing. Gene deletions were obtained by replacing the ORF through single step homologous recombination with an antibiotic resistance cassette derived by PCR from the plasmids pFA6a-hphNT1, pFA6a-natNT2, pFA6a-kanMX6, pFA6a-CmLEU2 or pYM-N35 (Bähler et al., 1998; Janke et al., 2004).

Fragments of *FIR1, SPA2* or *SKT5* were expressed as GST- or 6xHis-fusions in *E.coli* strains BL21 or BL21 Gold. Fragments for the GST-fusions were amplified from yeast genomic DNA using primers containing *Bam*HI/*Eco*RI restriction sites. The PCR products were fused in-frame behind GST on a pGex2T vector (GE Healthcare, Buckinghamshire, UK). 6xHis-tagged fragments were amplified from yeast genomic DNA using primers containing *Sfi*I restriction sites. The products were inserted into the pES plasmid, downstream and in frame of a 6xHis-tag.

### *In vivo* Split-Ubiquitin interaction analysis

Large scale Split–Ubiquitin assays were performed as described (Dunkler et al., 2012). A library of 540 different α-strains each expressing a different N_ub_ fusion were mated with a *P_MET17_FIRI*-CRU-expressing a-strain. Diploids were transferred as independent quadruplets on SD media lacking methionine and containing 1 mg/ml 5-FOA, and different concentrations of copper sulfate to adjust the expression of the N_ub_ fusions. For small-scale interaction analysis, a- and *α-*strains expressing N_ub_ or C_ub_ fusion constructs were mated. The diploid cells were spotted onto SD-FOA medium in four 10-fold serial dilutions starting from OD_600_=1. Growth was recorded at 30°C every day for 2 to 5 days.

### *In vitro* binding assays

#### E.coli extracts

An overnight culture of Hisx6-Spa2_1314-end_- or GST-Fir1_1-410_-expressing *E. coli* BL21 Gold (GE Healthcare, Freiburg, Germany) was diluted into 500 ml of SB medium and incubated at 37°C to an OD_600_ of 0.8. The cultures were chilled to 18°C and incubated overnight at 18°C in the presence of 1 mM IPTG. Cells were centrifuged at 5000 xg for 10 min, resuspended in PBS, and centrifuged at 4000 xg for 10 min. The cell pellet was stored at −80°C. BL21 strains expressing GST-Fir1_741-876_, 6xHis-SKT5, or 6xHis6-Skt5_146-550_ were treated as above except that protein expression occurred in SB/1 mM IPTG for 4 h at 37°C.

The cell pellets were washed once in PBS and resuspended in 5 ml PBS containing protease inhibitor cocktail (Roche Diagnostics, Penzberg, Germany) and 1 mg/ml lysozyme. After 45 min on ice, cells were lysed by 3×2 min sonication with a Bandelin Sonapuls HD 2070 (Reichmann Industrieservice, Hagen, Germany). Extracts were clarified by centrifugation at 40,000 g for 10 min at 4°C. 6xHis-Spa2_1314-end_ expressing strain BL21 was lysed in IMAC Buffer A (50 mM KH2PO4 pH8.0, 300 mM NaCl, 20 mM imidazole) by lysozyme treatment and sonication, followed by IMAC purification on an Äkta purifier chromatography system using a 5ml HisTrap Excel column (GE Healthcare, Freiburg, Germany). The column was washed by a linear imidazole gradient (20-70 mM), followed by elution in 200 mM imidazole. Eluted proteins were subsequently transferred on a PD10 column into PBS and concentrated. Enriched 6xHis-Spa2_1314-end_ was finally loaded onto a gel-filtration column (HiLoad 16/600 Superdex 200pg) equilibrated in 20 mM MES, 100 mM NaCl, pH 6.7. Peak fractions were pooled and used for the experiments.

#### Binding assay

All incubation steps were carried out under rotation at 4°C. GST or GST-tagged proteins were immobilized from *E. coli* extracts on 100 μl Glutathione–Sepharose beads in PBS (GE Healthcare, Freiburg, Germany). After incubation for 1h at 4°C with either *E.coli* extracts or purified proteins, the beads were washed three times, the bound material was eluted with GST elution buffer (50 mM Tris, 20 mM reduced glutathione), and subjected to SDS-PAGE followed by Coomassie Blue staining and Western Blot analysis using anti-His, or anti-GST antibodies (Sigma-Aldrich, Darmstadt, Germany).

#### Far-Western blot

Enriched GST- and GST-Fir1_1-410_ were transferred onto nitrocellulose after SDS PAGE. The membrane was blocked for one hour in TBST 2% milk powder (w/v), washed for five min in 10 ml TBST, and incubated for one hour at 4°C with extracts (20-fold diluted in TBST) from *E.coli* expressing no additional protein or 6xHis-Skt5_146-550_. After washing three times for five minutes with 10 ml TBST, the membrane was sequentially incubated at room temperature for one hour with anti-His and conjugated anti-mouse antibodies (Sigma-Aldrich, Darmstadt, Germany) (Wu et al., 2007).

### Fluorescence microscopy

Microscopic observations were performed with two different fluorescence microscopes. The Axio Observer spinning disc confocal microscope (Zeiss) is equipped with an Evolve512 electron-multiplying charge-coupled device camera (Photometrics), a Plan-Apochromat 63×/1.4 oil differential interference contrast (DIC) objective and a 100x/1.4 oil differential interference contrast (DIC) objective. Fluorescence was excited with 488 and 561-nm diode lasers (Zeiss) and detected with high efficiency filter sets 38 (GFP) and 45 (Cherry), respectively. Time lapse experiments were carried out at 30°C in a PeCon Incubator controlled by a PeCon TempModule S1. Operations were performed with the ZEN2.6 (2012) software package (Zeiss).

The DeltaVision system (GE Healthcare) is provided with an IX71 microscope (Olympus), a 100× UPlanSApo 100× 1.4 oil ∞/0.17/FN26.5 objective (Olympus), a steady-state heating chamber (Weather station by Prescision Control), and a Photofluor LM-75 halogen lamp. The standard live cell filter set (eGFP excitation 470/40, emission 525/5; dsRed: excitation 580/20, emission 630/60) was used for fluorescence microscopy and images were recorded with either the Cascade II 512 electron-multiplying charge-coupled device camera (Photometrics) or the CoolSNAP HQ^2^ High-Speed CCD camera (Photometrics). All operations were performed with the SoftWoRx 6.1.3. software (GE Healthcare).

For all microscopic analyses yeast cultures were grown overnight in SD medium, diluted in 3–4 ml fresh SD medium, and grown for 2–3 h at 30°C to mid-log phase. Exposure times were adapted to the respective GFP- and mCherry-labelled proteins to reduce bleaching and phototoxicity.

Standard time-lapse experiments with the ZEISS microscope were carried out with the 63x objective in a Sarstedt 1-well on cover glass II incubation chamber. 100 μl culture was pipetted onto the glass bottom of the chamber and covered with a slice of standard solid SD medium. Images were taken in intervals of 2 minutes and obtained with a series of seven Z-slices and a distance of 0.5 μm between adjacent z-slices.

Standard time lapse experiments at the DeltaVision microscope were performed with the 100x objective and the Cascade II 512 camera. 1 ml mid-log phase cell culture was carefully pelleted and resuspended in 50 μl medium. 3 μl of this suspension was transferred on custom-designed glass slides containing solid SD medium with 1.8% agarose and immobilized with a coverslip. Images were taken in intervals of 2 minutes and obtained with a series of seven z-slices and a distance of 0.5 μm between adjacent z-slices.

### Analysis of microscopic data

Image analysis and signal quantifications were carried out with the open source software platforms Image J for the data obtained with the DeltaVision microscope or Fiji for the data obtained with the ZEISS microscope (Schneider et al., 2012; Schindelin et al., 2012).

### Quantitative evaluation of time lapse experiments

For the quantification of bud neck signals in time lapse series of cells undergoing cytokinesis we generally employed yeast strains expressing two different fusion proteins. The protein of interest was translationally fused to GFP while fusions of either the septin Shs1 or the type II myosin heavy chain Myo1 served as marker proteins for the onset of cytokinesis. Sum projections of all relevant Z planes in both channels served as basis for all measurements. If not noted differently, circular regions of interest (ROI) of exactly 8 pixels diameter were selected covering the bud neck region (1, bud neck signal), a region without obvious signal within either the daughter or mother cytosol (2, cytosolic signal) and an area outside the cell (3, background). The mean grey value of ROI 1-3 in both channels at 19 timepoints covering the period of cytokinesis was measured. Data analysis was carried out with Microsoft Excel (Microsoft Excel for Mac, version 16.66.1) and GraphPad Prism (Prism 9 for MacOS, version 9.0.2). The background (3) was subtracted from values (1) and (2) in both channels at every timepoint and the resultant values (s1) were divided by the cytosolic readings (s2) to obtain the ratio of bud neck to cytosol signal for both channels at each time point (FU, fluorescent units). The Shs1-Cherry signal drops dramatically when the septin ring splits (defined as minute 18 in this study) while Myo1-Cherry contracts and disappears rapidly at minute 22. To facilitate temporal alignment of the readings originating from different cells each individual time series was normalized, meaning that the lowest value within each individual series was defined as 0% and the maximum value as 100% (RFU, relative fluorescent units). The RFU values obtained for the Cherry channel were plotted against time and utilized for a visual temporal alignment (facilitated by the loss/drop of the Myo1-/Shs1-Cherry signal) of the individual curves of this experiment. For each experiment the FU as well as the RFU series were aligned accordingly and the averages and standard deviations of each time point in the GFP as well as the Cherry channel were determined. Normalization reduces the quantitative deviation between individual curves, facilitates alignment and is useful for illustrating the relative timing of signals of comparable intensities. In most cases we preferred to show the absolute FU as this data set provides additional information on relative signal strength. We are aware that absolute fluorescence signal intensities vary considerably between experiments. The shown quantitative differences could be reproduced repeatedly and independently and represent the visual impression of the pictures very accurately.

### Statistical evaluation

For all experiments where the curves of collated protein variants or genotypes displayed reproducibly deviating FU values, the statistical significance of the difference at selected time points was determined in GraphPad Prism. Kruskal-Wallis Anova tests were used for multiple comparisons, whereas Mann-Whitney tests were employed for the comparison of two data sets.

### Images time series

For the illustration of bud neck signals, representative cells of the standard time lapse experiments were chosen and Max Z-projections were created in ImageJ or Fiji. Brightness and contrast settings of the GFP and Cherry channels were manually adjusted.

### Specifications of selected microscopic experiments

*Figure 1D; Figure S1A:* The pictures in Fig. 1D were obtained with the Deltavision microscope utilizing the 100x objective and the CoolSNAP HQ^2^ camera. In a short time-lapse experiment 20 Z sections at 0,2 μm spacing were taken every 2 minutes. Images of the GFP- and Cherry-channels were deconvoluted with the softWoRx 6.1.3 software and subsequently 3D projected in ImageJ. *Figure 2F:* Images shown in Fig. 2F originate from standard Deltavision time lapse experiments. A Max projection of the GFP channel was superimposed on single Z sections of the bright field channel. Profiles were measured in single Z sections of the same cells. A linear ROI with the width of 1 pixel was drawn centrally through budding cells from the mother to the daughter starting and ending outside the cells. The average of the extracellular values of the GFP channel was subtracted from each data point. The value for each data point was divided by the average of the extracellular values of the GFP channel. *Figure 4D:* The Images of the bud neck in Fig. 4D were obtained from a single cell at 2 different time points with the 100x objective at the Zeiss microscope. Overlays of Sum projections of the GFP and Cherry channel are shown. The corresponding profiles were measured in single Z sections of the same cell. A short linear ROI with the width of 1 pixel was drawn centrally through the bud neck from the mother to the daughter cell. The measurements of both channels were normalized. *Figure 4E:* Due to the patchy cytosolic distribution of Chs3-GFP a circular selection of 13 pixels diameter in the mother cell was used as intracellular reference. *Figure 6A:* Due to the high level of cytosolic Fir1-GFP-signal in *cdh1Δ-cells*, the standard procedure for quantitative timelapse analyses led to an underestimation of the bud neck signal. We thus decided to divide the measured bud neck signal by the reference value obtained outside the cell in wildtype and *cdh1Δ* rather than a cytosolic reference point. *Figure 6C:* Images originate from standard Zeiss time lapse experiments. A Sum projection of the GFP channel was superimposed on a single Z section of the bright field channel. Profiles were measured in single Z sections of the same cells. A linear ROI with the width of 1 pixel was drawn centrally from the mother to the daughter starting and ending outside the cells. The value for each data point was divided by the average of the extracellular values of the GFP channel.

### Online supplemental material

Fig. S1 shows deconvoluted images of the bud neck of cells expressing Fir1-GFP and Fir1_ΔSIM_-GFP, the complementation of *siz1*Δ-*cells*, and the statistical evaluation of the fluorescence intensity profiles of Fig. 1E. Fig. S2 shows the Split-Ub interaction array, the fluorescence intensity profiles of Spa2-GFP in different genetic backgrounds, the complementation of *spa2*Δ-cells, and the statistical evaluation of the fluorescence intensity profiles of Fig. 2D. Fig. S3 shows the far-western analysis of Fir1_750-876_ probed with Spa2_1324-end_. Fig. S4 shows the fluorescence intensity profiles of different bud neck localized proteins in the presence and absence of *FIR1*, and the statistical evaluation of the fluorescence intensity profiles of Figs. 4B, C, E. Fig. S5 shows the input fractions of Fig 5B. Fig. S6 shows the statistical evaluation of the fluorescence intensity profiles of Figs. 6A, D. Table S1 lists the Split-Ub discovered interaction partners of Fir1. Tables S2, S3, S4 list the yeast strains, constructs and primers used and created in this study.

## Acknowledgement

We thank N. Schmid and S. Timmermann for technical assistance. We thank J. Dohmen (University of Cologne, Germany) for yeast strains, plasmids and advice.

## Author contributions

Conceptualization: N. Johnsson, J. Müller. Investigation: J. Müller, M. Furlan, B. Grupp, D. Settele. Writing - original draft: N. Johnsson; Writing - review & editing: N. Johnsson, J. Müller, M. Furlan. Supervision: N. Johnsson. Funding acquisition: N. Johnsson.

