## Supplementary Information for "Transient septin sumoylation steers a Fir1-Skt5 protein complex between the split septin ring"

### Supplementary Figures

#### Figure S1

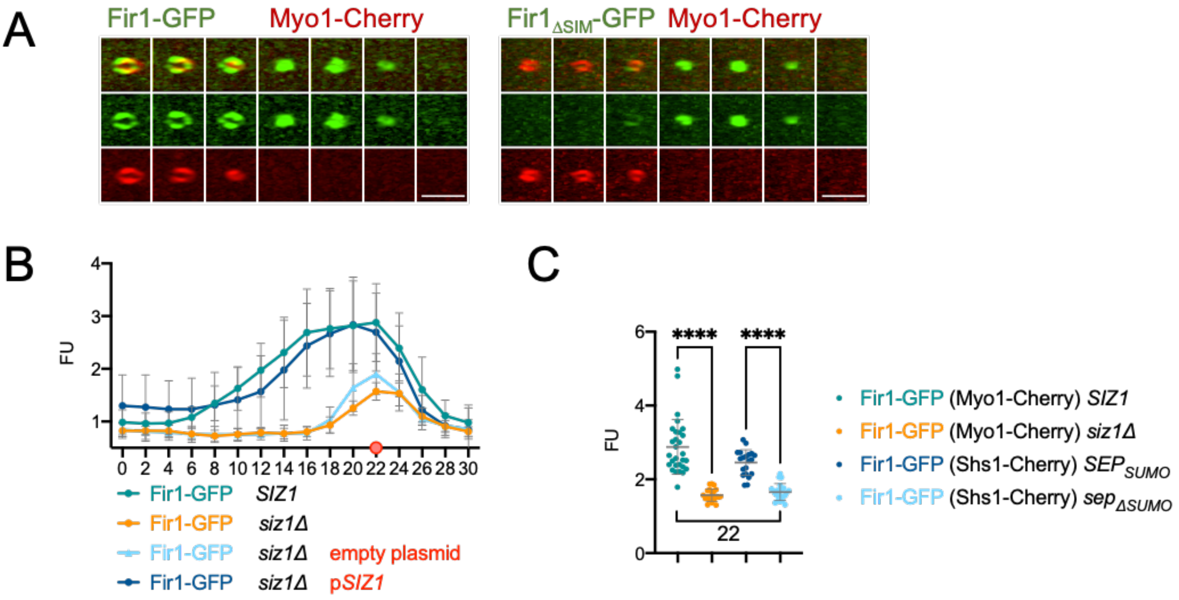


(A) Deconvoluted images of the bud neck of cells coexpressing Fir1-GFP (left panels) and Fir1$_{\Delta SIM}$-GFP (right panels) together with Myo1-Cherry, 6 minutes before and after complete Myo1 contraction. Overlay, GFP- and Cherry-channels are shown in top, middle, and lower panels. Scale bars = 3µm. (B) as in (Fig. 1C) but showing the complementation of *siz1*Δ-cells*.* Coexpression of plasmid-encoded *SIZ1* (p*SIZ1;* n=25) rescued Fir1-GFP signal timing and intensity, while the empty vector had no effect (empty plasmid; n=23). (C) Statistical evaluation of differences in signal intensities of Fir1-GFP in wildtype, *sep*$_{\Delta sumo}$ and *siz1*Δ cells at time point 22 of Fig. 1E. **** p<0,0001.

**Figure S2**

A

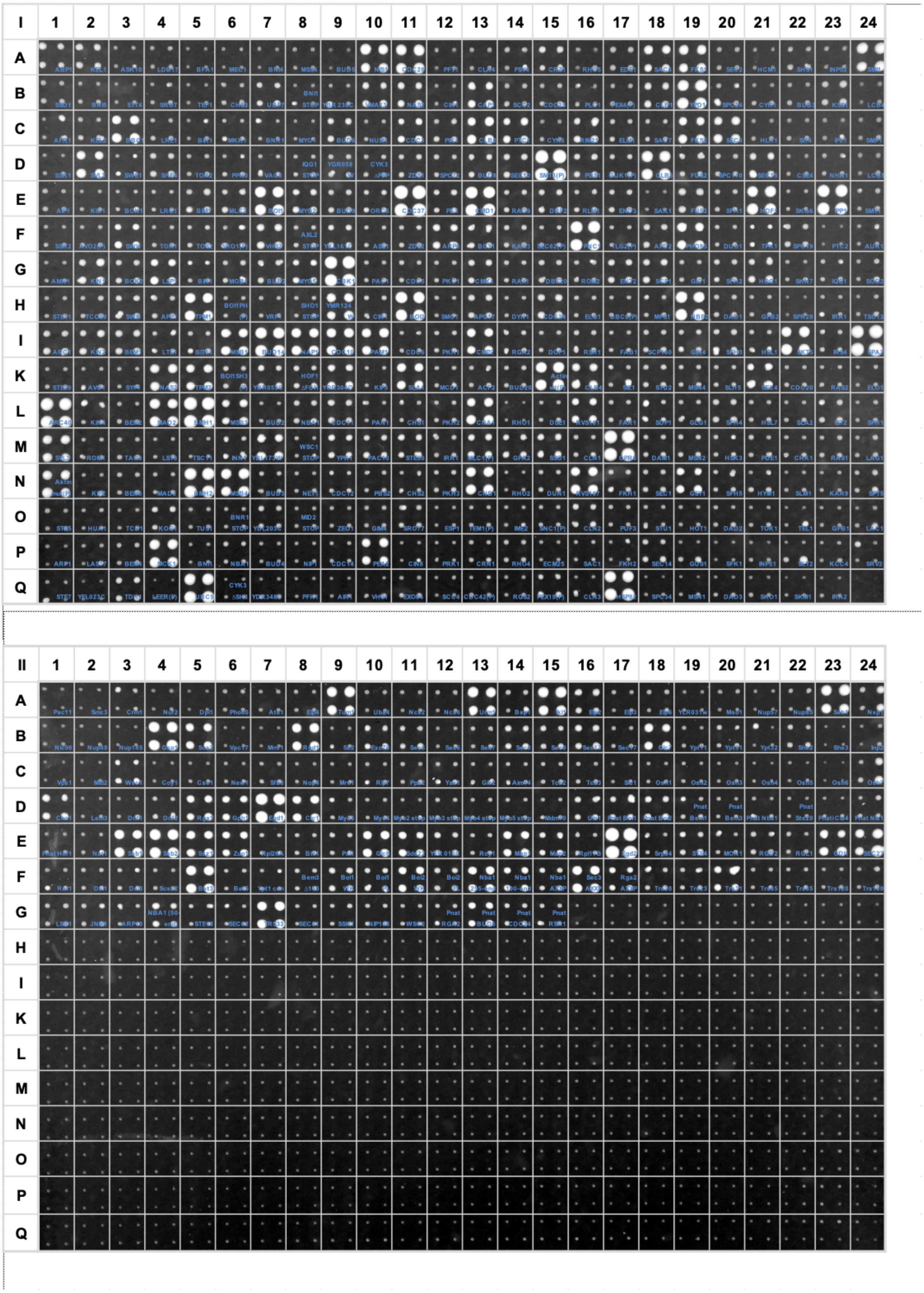

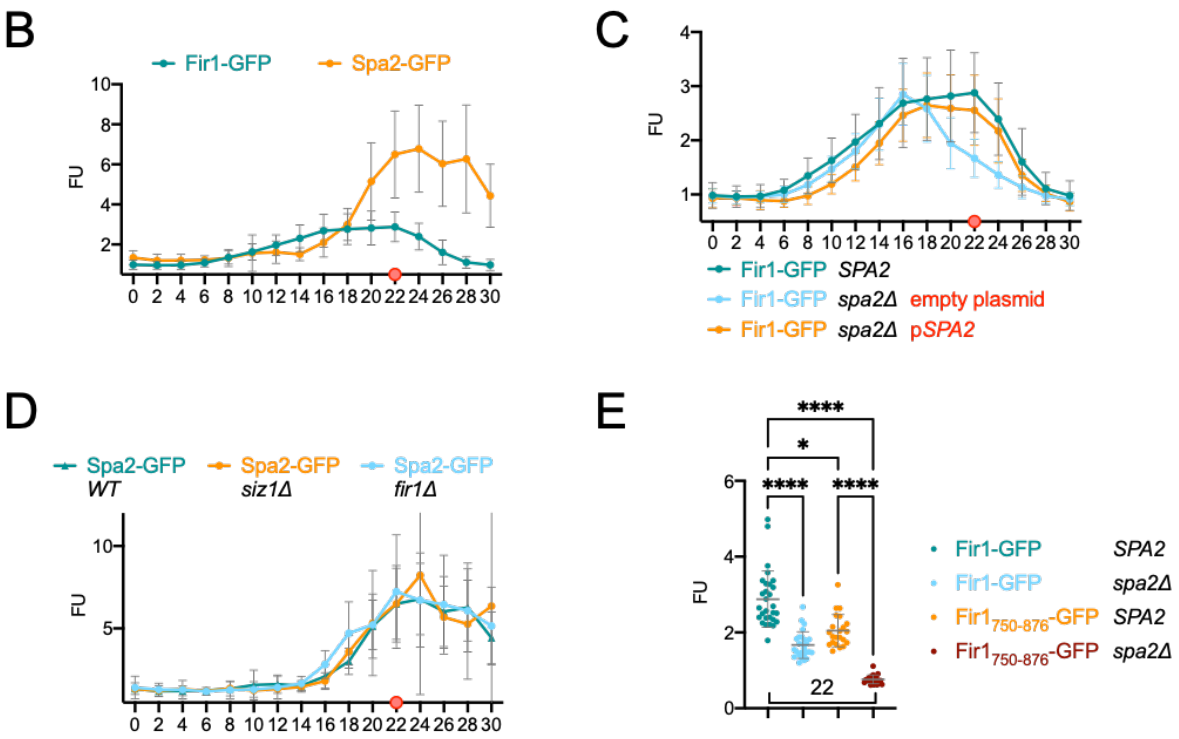


(A) Split-Ub interaction assay of 540 yeast strains coexpressing $P_{MET17}$-Fir1CRU each with a different $N_{ub}$ fusion protein. For every strain, cells of four independent matings were spotted as quadruplet on SD medium containing 5-FOA, 50 µM copper sulfate and 0 M methionine. Shown is the growth of the diploid yeast cells after 2 days at 30°C. Growth indicates protein-protein interaction. (B) As in (Fig. 1C) but with cells coexpressing Fir1-GFP, or Spa2-GFP with Myo1-Cherry. (C) As in (Fig. 1C) but showing the complementation of the reduction of the Fir1-GFP bud neck signal in *spa2*Δ-cells*.* Co-expression of plasmid-encoded *SPA2* (p*SPA2,* n=28) rescued timing and intensity of the Fir1-GFP signal, while the empty vector had no effect (empty plasmid, n=26). (E) As in (1C). The Spa2-GFP signal at the bud neck remains unaffected in timing and intensity by the deletion of *SIZ1* (*siz1*Δ; n=17*),* or *FIR1* (*fir1*Δ; n=13*).* (E) Statistical evaluation of differences in signal intensities at time point 22 of Fig. 2D between Fir1-GFP in wildtype- and *spa2*Δ-cells, between $Fir1_{750\text{-}876}$-GFP in wildtype- and *spa2*Δ-cells, and between Fir1-GFP an $Fir1_{750\text{-}876}$-GFP in wildtype- and *spa2*Δ-cells. **** p<0,0001; * p=0,0173.

**Figure S3**

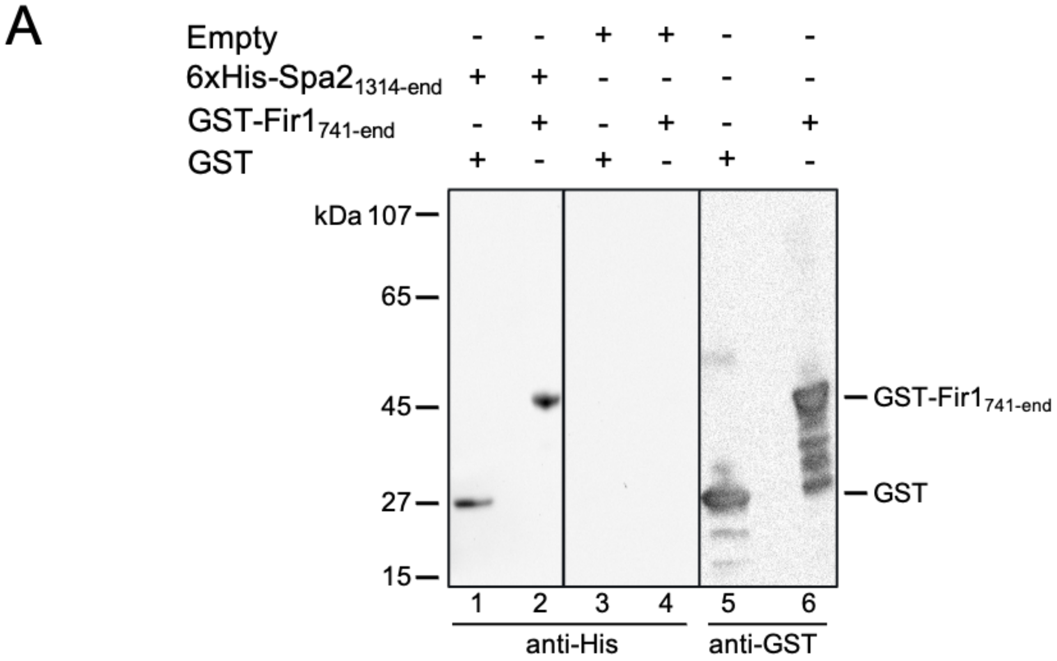


(A) Equal amounts of *E.coli* extracts containing GST (lanes 1, 3, 5), or GST-Fir1$_{741\text{-}876}$ (2, 4, 6) were separated by SDS-PAGE, transferred to nitrocellulose, and incubated with extracts from *E.coli* expressing no additional protein (lanes 3, 4), or 6xHis-Spa2$_{1324\text{-}end}$ (lanes 1, 2),. GST- or 6xHis-fusions were subsequently detected by anti-GST (lanes 5, 6), or anti-His antibodies (lanes 1-4).

**Figure S4**

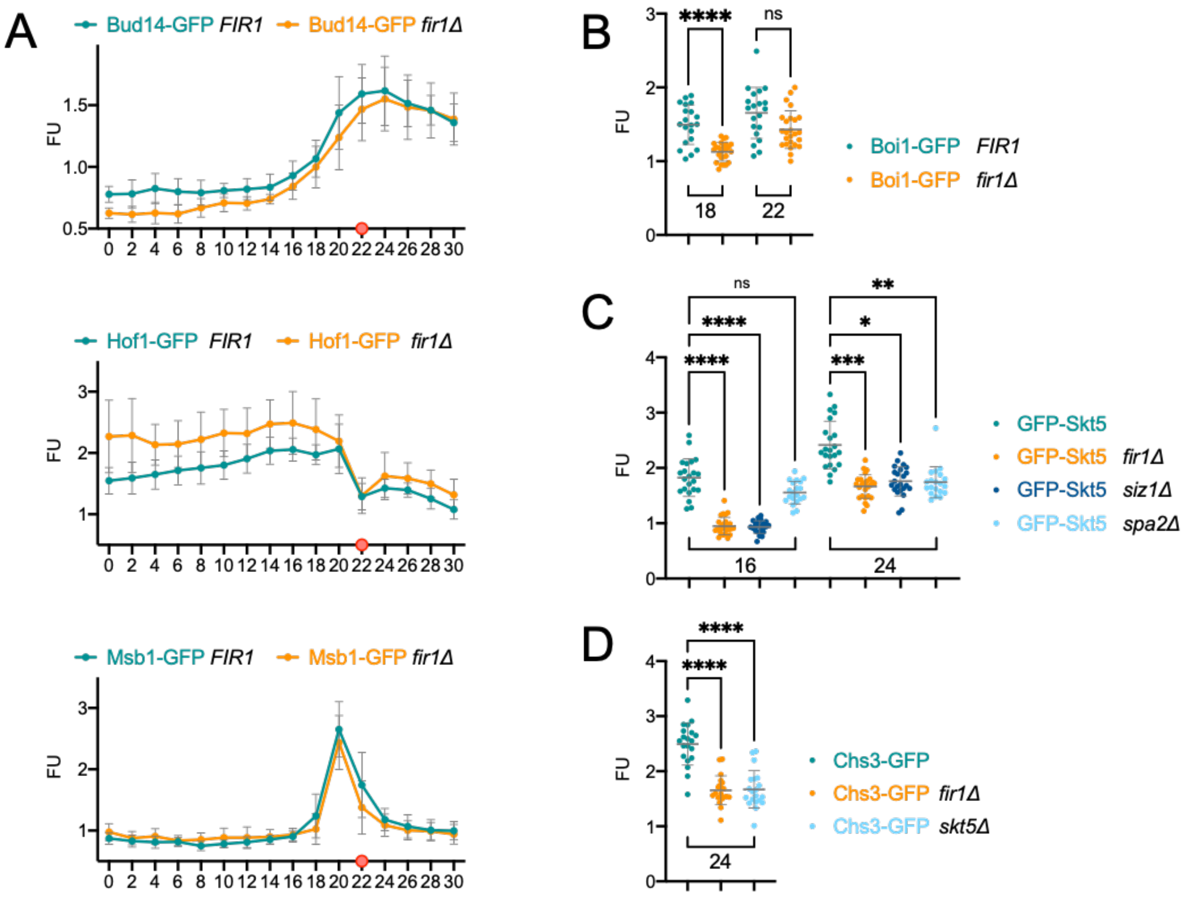


As in (1C) but with cells coexpressing Myo1-Cherry with Bud14-GFP (upper panel; widtype n=9; *fir1*Δ n=8), Hof1-GFP (middle panel; wildtype n=17; *fir1*Δ n=22), or Msb1-GFP (lower panel: wildtype n=13; *fir1*Δ n=10) with or without *FIR1.* (B) Statistical evaluation of differences in signal intensities between Boi1-GFP in the presence and absence of *FIR1* at time points 18 and 22 of Fig. 4B. **** p<0,0001; ns not significant. (C) Statistical evaluation of differences in signal intensities between GFP-Skt5 in wildtype cells and strains lacking *FIR1, SIZ1,* or *SPA2* at time points 16 and 24 of Fig. 4C. **** p<0,0001; *** p=0,0006; **=0,0039; ns not significant. (D) Statistical evaluation of differences in signal intensities between Chs3-GFP in the presence and absence of *FIR1*, or *SKT5* at time point 24 of Fig. 4E. **** p<0,0001.

**Figure S5**

A

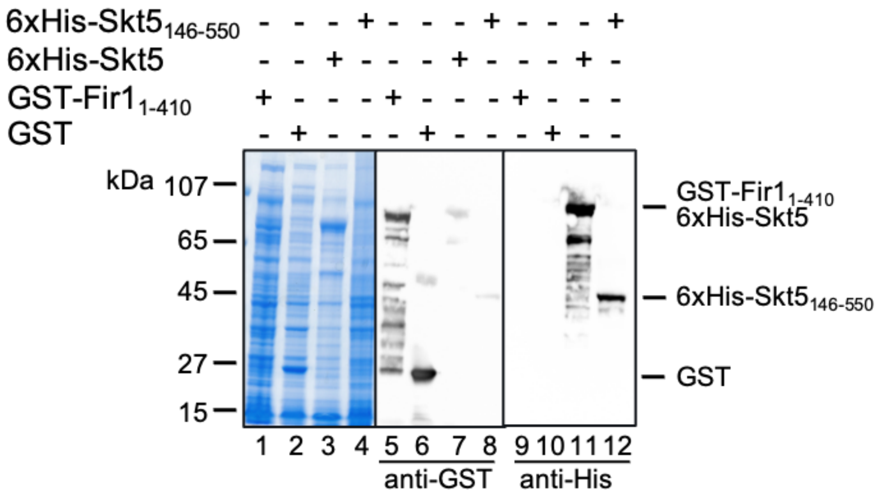


(A) Inputs for Fig 5B. *E. coli* extracts expressing GST-Fir1$_{1\text{-}410}$ (lanes 1, 5, 9), GST (lanes 2, 6, 10), 6xHis-Skt5 (lanes 3, 7, 11), or 6xHis-Skt5$_{146\text{-}550}$ (lanes 4, 8, 12) were separated by SDS-PAGE and either stained with Coomassie (lanes 1-4), or transferred to nitrocellulose and stained with anti-GST (lanes 5-8), or anti-His antibodies (lanes 9-12).

**Figure S6**

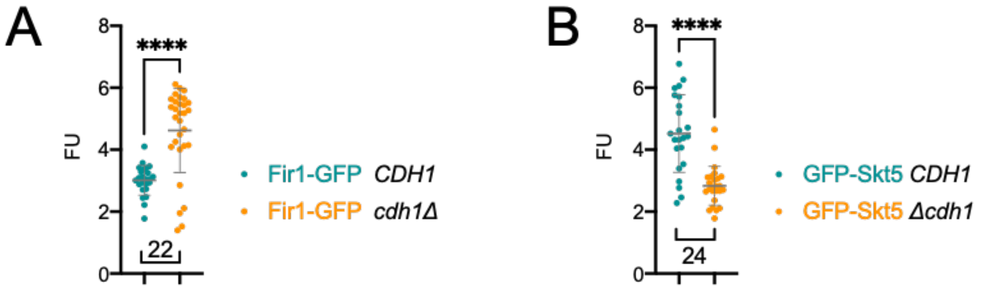


(A) Statistical evaluation of differences in signal intensities between Fir1-GFP in the presence and absence of *CDH1* at time point 22 of Fig.6A. **** p<0,0001. (B) Statistical evaluation of differences in signal intensities between GFP-Skt5 in in the presence and absence of *CDH1* at time point 24 of Fig. 6D. **** p<0,0001.

### Supplementary Tables

#### Table S1

Interaction partners of Fir1

| | Fir1CRU |
|---|---|
| $N_{ub}$ fusion proteins | Nis1 |
| | Sac6 |
| | Clb1 |
| | Clb2 |
| | Ptc1 |
| | Sec3 |
| | Smt3 |
| | Clb3 |
| | Boi1 |
| | Hof1 |
| | Cbk1 |
| | Epo1 |
| | Boi2 |
| | Cmp2 |
| | Bud14 |
| | Msb1 |
| | Skt5 |
| | Spa2 |
| | Clb4 |
| | Sla1 |
| | Cna1 |
| | Cnb1 |
| | Pea2 |
| | Ubc9 |
| | Glc7 |
| | Rga1 |
| | Rgd1 |
| | Gdi1 |
| | Sec3ΔPxxP |
| | Glc8 |
| | Msb4 |
| | Nbp2 |

List of $N_{ub}$ labeled interaction partners of Fir1CRU after subtraction of chaperones and false positives as defined by Hruby et al. (Hruby et al., 2011). Red letters indicate the newly identified interaction partners.

**Table S2**

List of *S. cerevisiae* strains used and created in this study.

| name | Relevant genotype | reference |
|---|---|---|
| JD47 | MATa, his3-Δ200, leu2-3, 112 lys2-801, trp1-Δ63, ura3-52 | Dohmen et al., 1995 |
| JD53 | MATα, his3-Δ200, leu2-3, 112 lys2-801, trp1-Δ63, ura3-52 | Dohmen et al., 1995 |
| EJY316 | *MATa cdc3-R4,11,30,63-HA::TRP1 cdc11-R412-HA::HIS3 shs1-R426,437-HA::HIS3* | Johnson and Blobel, 1999 |
| YJM_258 | JD47 pMet Fir1 ΔSIM CRU 303 | this manuscript |
| STY1401 | JD47 pMet Fir1 1-410 CRU 303 | this manuscript |
| STY1307 | JD47 pMet Fir1 CRU 303 | this manuscript |
| STY1555 | JD47 pMet Fir1Δ801-820 CRU 303 | this manuscript |
| YAD3517 | JD53 Nub Spa2 Δ281-428 | this manuscript |
| NS_1 | ULM53 Nub Spa2 | this manuscript |
| YAD3569 | ULM53 Nub Spa2 Δ1275-end | this manuscript |
| NS_2 | ULM53 Nub Spa2 Δ2-150 | this manuscript |
| NS_3 | ULM53 Nub Spa2 Δ2-188 | this manuscript |
| YJM271 | JD47 Fir1-GFP, Myo1-Cherry 306 | this manuscript |
| YJM314 | JD47 Fir1 750-876-GFP, Myo1-Cherry 306 | this manuscript |
| YJM387 | JD47 Fir1-GFP, Shs1-Cherry 306 | this manuscript |
| YJM381 | JD47 Fir1Δ801-820-GFP, Myo1-Cherry 306 | this manuscript |
| YJM281 | JD47 Fir1ΔSIM-GFP, Myo1-Cherry 306 | this manuscript |
| YJM383 | JD47 Fir1ΔSIM, Δ801-820-GFP, Myo1-Cherry 306 | this manuscript |
| YJM430 | EJY316 Fir1-GFP(HpH), Shs1-Cherry 306 | this manuscript |
| YAD4181 | JD47 cdh1::HpH, Fir1-GFP 304, Myo1-Cherry 306 | this manuscript |
| YAD4181 | JD47 cdh1::HpH, Fir1-GFP 304, Myo1-Cherry 306 | this manuscript |
| YJM405 | JD47 siz1::HpH Fir1-GFP, Myo1-Cherry 306 | this manuscript |
| YAD4196 | JD47 Spa2 1-1274, Fir1-GFP 304, Myo1mCherry 306 | this manuscript |
| YJM306 | JD47 spa2::HpH, Fir1-GFP, Myo1-Cherry 306 | this manuscript |
| YJM307 | JD47 spa2::HpH, Fir1ΔSIM-GFP, Myo1-Cherry 306 | this manuscript |
| YJM380 | ULM53 spa2::HpH, Fir1 750-876-GFP, Myo1-Cherry 306 | this manuscript |
| YAD4093 | JD47 Spa2-GFP 304, Myo1-Cherry 306 | this manuscript |
| YJM409 | JD47 fir1::HpH Spa2-GFP 304, Myo1-Cherry 306 | this manuscript |
| YJM411 | JD47 siz1::HpH Spa2-GFP 304, Myo1-Cherry 306 | this manuscript |
| YAD4198 | JD47 Spa2 1-1274-GFP 304, Myo1-Cherry 306 | this manuscript |
| YJM441 | JD47 GFP-Skt5, Myo1-Cherry 306 | this manuscript |
| YJM453 | JD47 GFP-Skt5, Shs1-Cherry 306 | this manuscript |
| YJM469 | JD47 GFP-Skt5, Shs1-Cherry 306 | this manuscript |
| YAD4160 | JD 47 spa2::HpH GFP-SKT5, Myo1-Cherry | this manuscript |
| YJM465 | JD47 cdh1::HpH GFP-Skt5, Myo1-Cherry 306 | this manuscript |
| YJM467 | JD47 cdh1::HpH GFP-Skt5, Shs1-Cherry 306 | this manuscript |

| | | |
|---|---|---|
| YJM443 | JD47 fir1::HpH GFP-Skt5, Myo1-Cherry 306 | this manuscript |
| STY1586 | JD47 siz1::HpH, GFP-Skt5, Myo1 Cherry 306 | this manuscript |
| YAD4162 | JD47 Cbk1-GFP 306, Myo1-Cherry 304 | this manuscript |
| YAD4164 | JD47 fir1::HpH Cbk1-GFP 306, Myo1-Cherry 304 | this manuscript |
| YAD4229 | JD47 Chs3-GFP 304, Myo1-Cherry 306 | this manuscript |
| YAD4230 | JD47 fir1::HpH, Chs3 GFP 304, Myo1-Cherry 306 | this manuscript |
| YAD4231 | JD47 Skt5::HpH, Chs3-GFP 304- Myo1-Cherry 306 | this manuscript |
| YJM146 | JD47 Boi1-GFP 304, Myo1-Cherry 306 | this manuscript |
| YJM449 | JD47 fir1::HpH, Boi1-GFP 304, Myo1-Cherry 306 | this manuscript |
| YJM354 | JD 47 Hof1-GFP 304, Myo1-Cherry 306 | this manuscript |
| YJM355 | JD47 fir1::HpH Hof1-GFP 304, Myo1-Cherry 306 | this manuscript |
| STY1574 | JD47 Bud14-GFP 304, Myo1-Cherry 306 | this manuscript |
| STY1563 | JD47 fir1::HpH, Bud14-GFP 304, Myo1-Cherry 306 | this manuscript |
| STY1576 | JD47 Msb1-GFP 304, Myo1-Cherry 306 | this manuscript |
| STY1564 | JD47 fir1::HpH, Msb1-GFP 304, Myo1-Cherry 306 | this manuscript |

**Table S3**

List of constructed plasmids in this study.

| name | description | Ref |
|---|---|---|
| pML107 | *3'sgRNA, Cas9, AmpR, LEU2* | Laughery et al.,2015 |
| pFA6a hphNT1 | *hphNT1, AmpR* | Janke et al., 2004 |
| pFA6a natNT2 | *natNT2, AmpR* | Janke et al., 2004 |
| pFA6a kanMX6 | *kanMX6, AmpR* | Janke et al., 2004 |
| pFA6a CmLEU2 | *CmLEU2, AmpR* | Janke et al., 2004 |
| pYM-N35 | *pMet, natNT2, AmpR* | Janke et al., 2004 |
| PYM25_GFPS65T (UJ602) | *GFPS65T hphNT1, AmpR* | this manuscript |
| PYM26_GFPS65T (UJ603) | *GFPS65T klTRP1, AmpR* | this manuscript |
| pGADT7 delta BamHI (UJ267) | *Amp, LEU2* | Clontech |
| pML107 Fir1_96 (UJ609) | *FIR1-sgRNA (PAM site 96), Cas9, AmpR, LEU2* | this manuscript |
| pML107 Fir1_2436 (UJ618) | *FIR1-sgRNA (PAM site 2436), Cas9, AmpR, LEU2* | this manuscript |
| pML107 Fir1_2271 (UJ591) | *FIR1-sgRNA (PAM site 2271), Cas9, AmpR, LEU2* | this manuscript |
| pML107 Skt5_17 (UJ624) | *SKT5-sgRNA (PAM site 17), Cas9, AmpR, LEU2* | this manuscript |
| pML104 Spa2-4029 (#1624) | *SPA2-sgRNA (PAM site 4029), Cas9, AmpR, URA3* | this manuscript |
| Fir1 (nt2177-end)-CRU 303 (#3733) | Fir1-CRU, Amp, His3 | this manuscript |
| Fir1 1-410 CRU 303 (#3754) | Fir1 1-410 -CRU, Amp, His3 | this manuscript |
| 6xHis-Spa2 1314-end (#1915) | pES Spa2 c-term (AS1314-End), Amp | this manuscript |

| | | |
|---|---|---|
| pGex-2T Fir1 741-end (#1946) | $P_{tac}$-GST Fir1 741-end, $Amp^R$ | this manuscript |
| pGex-2T Fir1 1-410 (#2029) | $P_{tac}$-GST Fir1 1-410, $Amp^R$ | this manuscript |
| 6xHis-Skt5 (#2066) | pES Skt5, AmpR | this manuscript |
| 6xHis-Skt5 146-550 (#2067) | pES Skt5 (146-550), Amp | this manuscript |
| Shs1-Cherry 306 | *Shs1-Cherry, AmpR, URA3* | this manuscript |
| Myo1-Cherry 304 | Myo1-Cherry, ampR, TRP1 | this manuscript |
| Myo1-Cherry 306 | *Myo1-Cherry, AmpR, URA3* | this manuscript |
| Fir1short (nt2461-end)-CRU 303 (UJ619) | Fir1short (nt2461-end)-CRU, Amp, His3 | this manuscript |
| Bud14-GFP 304 (#172) | Bud14-GFP, Amp, TRP1 | this manuscript |
| Spa2-GFP 304 (#362) | Spa2-GFP, Amp, TRP1 | this manuscript |
| Cbk1-GFP 306 (#460) | Cbk1-GFP, ampR, URA3 | this manuscript |
| Boi1-GFP 304 (#476) | Boi1-GFP, Amp, TRP1 | this manuscript |
| Msb1-GFP 304 (#1792) | Msb1-GFP, Amp, TRP1 | this manuscript |
| Fir1(nt2177-end)-GFP 304 (#3735) | Fir1-GFP, Amp, TRP1 | this manuscript |
| Spa2 ∆1275-end-GFP 304 | Spa2 ∆1275-end-GFP, Amp, His3 | this manuscript |
| Chs3-GFP 304 (#4072) | Chs3-GFP, Amp, TRP1 | this manuscript |
| Hof1-GFP 304 (UJ547) | Hof1-GFP, Amp, TRP1 | this manuscript |
| pGADT7 PSpa2-Spa2 (UJ612) | *SPA2, Amp, LEU2* | this manuscript |
| Siz1 315 (UJ620) | *SIZ1, Amp, LEU2* | this manuscript |

**Table S4**

List of primers used and created in this study.

| name | description | sequence |
|---|---|---|
| JMn266 | do Fir1∆SIM | TAATACCATTTACGAAAAAAGAGATGGTAAAATGGTTGAGGTAGG CCTCAAAAATAACGATATATCACGAACAAGAGTCT |
| JMn310 | do Fir1 ATG 750 | GCAATTAGATTGCGCTGGGATAGATCTGAAGCTTTTTTTCCGATT **ATG**GAAAAAAGAGATGGTAAAATGGTTGAGGTTATTCTTTTAGAT GAG |
| JMn325 | do Fir1 ∆801-820 | GCTAAAAATGAGCAACAGAAGAAACGTCTAAGCCATTGCAACGAA AATGAGAAGCATAAATTCTCAGAAAAAGGACAACAAACAAAGCCA |
| #6329 | doSpa2 1-1274 | AAACATTATAGTGATGATGATGATTCTAGCTATCAATTTGTTCCCT AAAGTTAGTAGTATCTAGAGGAGAAAGAAAAGGAAGACAAAGAC |
